# Spatial tumor microenvironmental architecture of chemoradiotherapy-resistant residual esophageal squamous cell carcinoma

**DOI:** 10.64898/2026.06.01.729308

**Authors:** Sungwoo Bae, Jungyoon Ohn, Hongyoon Choi, Hye Seung Lee, Bhumsuk Keam, Chang Hyun Kang, Hak Jae Kim, Kwon Joong Na, Byoung Hyuck Kim

**Author notes:** **Correspondence and reprint requests:** Byoung Hyuck Kim, MD, PhD, Department of Radiation Oncology, Seoul National University College of Medicine, Seoul Metropolitan Government – Seoul National University Boramae Medical Center, Seoul, Republic of Korea (07061), Kwon Joong Na, MD, PhD, Portrai, Inc, Seoul, Republic of Korea (04798); Department of Thoracic and Cardiovascular Surgery, Seoul National University Hospital, Seoul National University College of Medicine, Seoul, Republic of Korea (03080).

## Abstract

**Purpose:** Residual ESCC after neoadjuvant CCRT reflects incomplete treatment response and carries a high risk of recurrence. We sought to define the spatial tumor microenvironmental features of CCRT-resistant residual ESCC — using tumor regression grade (TRG) 2–3 tumors as a model of poor pathologic response — with mechanistic refinement at single-cell resolution and outcome assessment in internal and external cohorts.

**Experimental Design:** We profiled post-CCRT ESCC tissue microarray (TMA) cores by Visium FFPE whole-transcriptome spatial transcriptomics and orthogonal Xenium single-cell-resolution in situ profiling on serial sections from the same TMA blocks. Spot-level cell-type composition was inferred by CellDART deconvolution against a published ESCC single-cell RNA sequencing (scRNAseq). Local malignant-cell-enriched regions were defined by CancerFinder. We additionally re-analyzed the scRNAseq dataset to characterize SPP1⁺ and CXCL5⁺ macrophage states and their ligand-receptor signaling using CellChat. Outcome associations were evaluated in the SNUH cohort and externally tested in the TCGA ESCC cohort.

**Results:** TRG 2–3 residual tumors exhibited effector immune-cell exclusion from malignant-cell-enriched regions, SPP1⁺/CXCL5⁺ macrophage accumulation, microvascular rarefaction, and proliferative-metabolic reprogramming. Single-cell re-analysis confirmed that CXCL5⁺ macrophages constitute a transcriptional subset of the broader SPP1⁺ macrophage population. CellChat showed that SPP1⁺ macrophages dominantly signal through SPP1–CD44 and SPP1–integrin axes to stromal and epithelial targets, and additionally engage immunosuppressive NECTIN2–TIGIT, CD86–CTLA4, and LGALS9–HAVCR2 programs with CD8 T cells, providing a mechanistic context for local immune exclusion. Visium LIANA and Xenium distance-gradient profiling localized SPP1-associated signaling preferentially to tumor cells and CAFs. Higher SPP1 expression and lower endothelial abundance were associated with shorter disease-free survival in the SNUH cohort. Higher SPP1 expression was also associated with shorter disease-free survival in the independent TCGA ESCC cohort.

**Conclusions:** CCRT-resistant residual ESCC is characterized by a spatially organized tumor microenvironmental niche centered on SPP1-associated macrophage programs and microvascular rarefaction. These spatially resolved findings identify candidate macrophage- and vasculature-targeted axes for overcoming treatment resistance.

**Translational Relevance:** Patients with esophageal squamous cell carcinoma (ESCC) who harbor residual disease after neoadjuvant chemoradiotherapy (CCRT) remain at high risk of recurrence, yet the spatial biology of this resistant residual state has been poorly defined. Here, we combine Visium whole-transcriptome and Xenium single-cell-resolution spatial transcriptomics with single-cell RNA-seq re-analysis and external TCGA context to define the tumor microenvironmental architecture of CCRT-resistant residual ESCC. The resistant niche is characterized by effector immune-cell exclusion from malignant-cell-enriched regions, accumulation of SPP1⁺/CXCL5⁺ macrophage programs that signal to tumor cells and cancer-associated fibroblasts through SPP1–CD44 and SPP1–integrin axes, microvascular rarefaction, and proliferative-metabolic reprogramming. These findings define a coordinated spatial tumor microenvironmental signature of CCRT-resistant residual ESCC and nominate macrophage–tumor/stromal interactions and vascular injury as biologic axes that may help guide future strategies to overcome treatment resistance.

## INTRODUCTION

Locally advanced esophageal squamous cell carcinoma (ESCC) is commonly treated with neoadjuvant chemoradiotherapy (CCRT) followed by surgical resection (1–3). Although a subset of patients achieves a pathologic complete response, a substantial proportion harbor residual disease at the time of surgery. This residual post-CCRT state is clinically important because poor pathologic response, particularly TRG 2–3 disease, is associated with a higher risk of recurrence and unfavorable long-term outcomes (6, 7). With the recent integration of adjuvant immunotherapy after neoadjuvant CCRT and surgery into clinical practice, there is a growing need to understand not only how much tumor remains after CCRT, but also which biologic features characterize the residual treatment-resistant state (4, 5).

Resistance to CCRT is unlikely to be explained by malignant-cell-intrinsic programs alone. Residual tumor cells persist within a reorganized tumor microenvironment (TME) shaped by prior chemoradiotherapy, immune remodeling, stromal responses, and vascular injury or recovery (8–10). Bulk transcriptomic approaches cannot resolve how immune, macrophage, fibroblast, endothelial, and malignant-cell programs are positioned relative to one another within residual tissue. Spatial transcriptomic (ST) technologies provide an opportunity to address this problem directly by defining the local architecture of treatment-resistant niches (11, 12).

Here, we integrated Visium FFPE whole-transcriptome spatial transcriptomics with Xenium single-cell-resolution in situ profiling to decipher spatial TME features associated with poor pathologic response in post-CCRT ESCC. We focused on patient-level comparisons between TRG 0–1 and TRG 2–3 tumors, on the spatial positioning of cellular states relative to malignant-cell-enriched regions, and on orthogonal single-cell-resolution refinement of key cellular interactions, complemented by external single-cell RNA-seq and TCGA-ESCA contexts. Finally, because resistant TME features may carry clinical implications, we asked whether emergent niche features were associated with disease-free survival as supportive evidence of their translational relevance.

## MATERIALS AND METHODS

### Ethics statement

This retrospective study was approved by the Institutional Review Board of Seoul National University Hospital and was conducted in accordance with the Declaration of Helsinki. Because archival FFPE tissue and retrospective clinical data were used, the requirement for written informed consent was waived by the Institutional Review Board.

### Patient cohort and tissue specimens

Patients with histologically confirmed ESCC who received neoadjuvant CCRT followed by curative-intent surgical resection at Seoul National University Hospital were retrospectively identified. Pathologic response was assessed on resection specimens by board-certified pathologists using an institutional 0–3 TRG scale consistent with prior ESCC reports (6, 7). Patients were grouped as good responders (TRG 0–1) or poor responders/CCRT-resistant residual disease (TRG 2–3). Tissue microarrays (TMAs) were constructed using 2-mm cores sampled from representative tumor-containing or tumor-bed regions, with one core per patient to avoid patient-level pseudo-replication. This design allowed patient-level comparison of local post-CCRT tissue architecture while acknowledging that each core represents a sampled region rather than the entire residual tumor bed.

### Visium FFPE spatial transcriptomics and analysis

Visium Spatial Gene Expression for FFPE assays (10x Genomics, Pleasanton, CA, USA) were performed across five TMA slides containing 56 evaluable post-CCRT ESCC cores. Raw data were processed using Space Ranger (version 3.1.3.). Downstream analyses were performed in Python using scanpy/anndata-based workflows together with additional spatial and statistical analysis packages. To integrate spots across TMA slides while preserving spatial adjacency, we used STAligner (13). Spot-level cell-type composition was inferred using CellDART (14) with a published ESCC single-cell RNA-seq atlas as reference (15). Cell-state labels defined in the original GSE221561 publication were retained without reclustering, yielding 46 fine-grained cell states. Reference marker distributions relevant to macrophage and endothelial annotations are summarized in **Supplementary Figure S1**.

To annotate local malignant-cell-enriched regions within each core, we used CancerFinder, a deep-learning-based method that infers malignant spots from transcriptomic profiles (16). Briefly, CancerFinder applies a pretrained spot-level classifier to assign each Visium spot a tumor versus non-tumor label according to transcriptomic features representative of malignant epithelium. The resulting binary tumor mask was used to define local malignant-cell-enriched boundaries within each sampled TMA core. Concordance between CancerFinder output and pathologist assessment is shown in **Supplementary Figure S2**.

For each Visium spot, we calculated the signed distance to the nearest boundary of the CancerFinder-defined malignant-cell-enriched region. Negative values indicate spots within malignant-cell-enriched regions, positive values indicate surrounding nonmalignant tissue within the same TMA core, and zero denotes the local boundary. Spots were binned at 100-µm intervals. Because these analyses were performed on 2-mm TMA cores, signed distance should be interpreted as local spatial context within sampled tumor-containing regions rather than as whole-tumor architecture. This approach was used to determine whether key immune, macrophage, stromal, and endothelial programs differed not only in abundance but also in their spatial positioning relative to malignant-cell-enriched regions.

### Xenium in situ spatial transcriptomics

Xenium in situ profiling was performed on serial sections from the same TMA blocks using the predesigned Xenium Human Multi-Tissue and Cancer panel (480 genes). Cell segmentation was performed using Proseg (17), yielding 882,604 segmented cells across 56 cores, of which 50 were matched to evaluable Visium cores at the core level. Cell-type annotation used ResolVI-based denoising/correction together with TACCO reference transfer (18) based on the same published ESCC single-cell atlas (15), producing major cell-type labels including Tumor_Cyc, Tumor_Diff, Cytotoxic_T_NK, TAM_DC, CAF, vSMC, Endothelial, Plasma, and Neutrophils.

Because Xenium was generated from serial sections of the same TMA blocks used for Visium, Xenium was treated as an orthogonal high-resolution refinement layer rather than as an independent validation cohort. Its primary role was to separate receiver cell types at single-cell resolution, refine the localization of macrophage-rich niches, and audit candidate SPP1-associated receiver programs. For receiver analyses, SPP1-high macrophages were defined as TAM_DC cells with SPP1 expression at or above the dataset median, and the distance from each Xenium cell to the nearest SPP1-high macrophage was calculated using cKDTree.

### Pathway activity inference

Pathway analyses were performed on Visium whole-transcriptome data only. Spot-level pathway activities were inferred using decoupleR (19). PROGENy pathway scores were estimated using multivariate linear modeling with pathway-responsive gene weights (20), and MSigDB HALLMARK gene sets were summarized using over-representation analysis. For patient-level analyses, pathway activities were aggregated across spots within each core, and TRG 0–1 versus TRG 2–3 differences were assessed using the Mann–Whitney U test with Benjamini–Hochberg correction within each pathway family.

### Ligand–receptor interaction analysis

Ligand–receptor interactions were inferred from Visium data using LIANA (21). Per-core ligand–receptor patterns were summarized after non-negative matrix factorization within tumor-enriched and stromal spatial domains. We specifically examined SPP1-containing and ITGB1-related ligand–receptor pairs because macrophage-associated SPP1 programs emerged as a prominent feature of the TRG 2–3 resistant niche. Spatial overlap analyses were performed using STopover (22). Top SPP1-containing and ITGB1-containing ligand–receptor pairs are summarized in **Supplementary Tables S2** and **S3**, respectively.

### Single-cell re-analysis in a public ESCC scRNA-seq dataset

To dissect the SPP1⁺ and CXCL5⁺ macrophage states observed in the spatial data at single-cell resolution, we re-analyzed the published ESCC scRNA-seq atlas (15) using Seurat (v5). Cells were quality-filtered (200 < nFeature_RNA < 2,500; percent mitochondrial < 15%), log-normalized (scale factor 10,000), and 2,000 highly variable features were selected for principal component analysis followed by graph-based clustering (FindNeighbors/FindClusters; resolution 0.3 on the first 20 principal components) and UMAP embedding. Cluster-defining markers were identified by Wilcoxon rank-sum test (FindAllMarkers; only.pos = TRUE; min.pct = 0.25; logfc.threshold = 0.25), and 21 cell types were annotated based on canonical markers.

To resolve macrophage states, the monocyte/macrophage compartment was further subclustered (FindSubCluster; resolution 0.8) and labeled based on SPP1 and CXCL5 expression patterns. Subclusters with predominant SPP1 expression but minimal CXCL5 expression were labeled SPP1⁺ macrophages; the subcluster with predominant CXCL5 expression (which also expressed SPP1) was labeled CXCL5⁺ macrophages; remaining myeloid subclusters were collectively labeled Monocyte_other.

To infer cell–cell communication, we constructed a CellChat object (CellChatDB.human; secreted, ECM, and contact pathways) (23) using the log-normalized RNA assay and final cell-type labels. After identification of overexpressed ligand-receptor pairs and overexpressed interactions per cell type, communication probabilities were computed (raw.use = TRUE), weak interactions were filtered (min.cells = 10), and pathway-level communication networks were aggregated. Subsequent visualization focused on signals originating from SPP1⁺ and CXCL5⁺ macrophages directed to all annotated target populations and on the SPP1, CXCL, and immune-checkpoint pathway families. Because this re-analysis used the same atlas as the spatial deconvolution reference, it is interpreted as single-cell-resolution mechanistic dissection of the spatial findings rather than as independent dataset validation.

### External validation in the TCGA ESCC cohort

To externally test prognostic associations of SPP1 expression, we used the TCGA Esophageal Carcinoma cohort (TCGA-ESCA, PanCancer Atlas 2018), restricted to histologically squamous cases. mRNA expression (RNA-seq V2 RSEM) and clinical data, including overall survival (OS) and disease-free survival (DFS), were merged at the patient level. Patients with non-positive follow-up time or missing event/time information were excluded, yielding 63 evaluable ESCC patients. Patients were dichotomized into high and low SPP1 expression groups using a median split. Survival was estimated by the Kaplan–Meier method and compared by the log-rank test.

### Outcome association analysis (SNUH cohort)

Outcome analyses in the SNUH cohort were performed to assess the clinical relevance of resistant-niche features that emerged from the spatial analyses. DFS was the primary endpoint. Locoregional recurrence-free survival (LRFS), distant metastasis-free survival (DMFS), and OS were considered secondary or exploratory endpoints. Time origin was the date of surgical resection. Continuous z-scored predictors were evaluated using Cox proportional hazards models, and Kaplan–Meier curves based on median splits were used for descriptive visualization only. Multivariable analyses were limited to parsimonious models consistent with the number of events.

### Statistical analysis

All primary patient-level analyses used one core per patient. Group comparisons used the Mann–Whitney U test with Benjamini–Hochberg correction within logical families, including cell states, pathways, and ligand–receptor pairs. Effect sizes are reported as rank-biserial correlation where appropriate. Spearman correlations were used for spatial gradient analyses, and one-sample Wilcoxon tests were used to evaluate the distribution of per-patient correlations. Two-sided P values < 0.05 were considered statistically significant.

## RESULTS

### A patient-level spatial atlas of post-CCRT ESCC

We profiled 56 post-CCRT ESCC patients. Selected clinicopathologic characteristics of the Visium cohort are summarized in **Supplementary Table S1**. The cohort comprised TRG 0 (n = 23), TRG 1 (n = 12), TRG 2 (n = 17), and TRG 3 (n = 4) tumors; the primary comparison contrasted good responders (TRG 0–1, n = 35) with poor responders (TRG 2–3, n = 21) (**Figure 1A**). We profiled 56 post-CCRT ESCC patients with Visium FFPE whole-transcriptome ST across five TMA slides (23,032 spots), together with 56 serial-section Xenium cores (882,604 cells), of which 50 were matched to evaluable Visium cores (**Figure 1A**). STAligner integration quality control is shown in **Supplementary Figure S3**.

**Figure 1.**
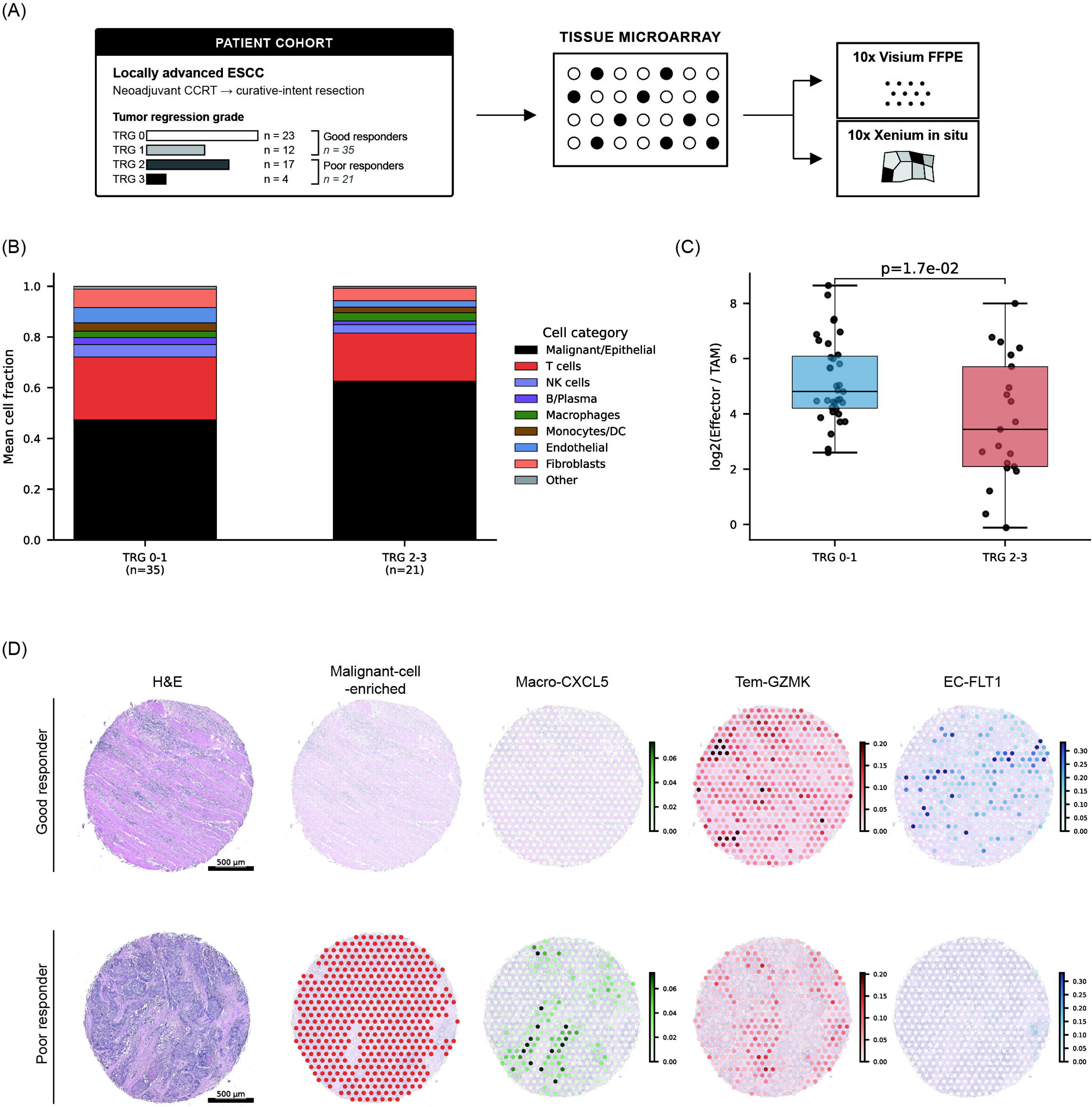
Cohort overview and patient-level spatial atlas of post-CCRT ESCC. (A) Study design schematic. Post-CCRT ESCC resection specimens from the SNUH cohort were assembled into tissue microarrays, followed by Visium FFPE and Xenium in situ spatial transcriptomic profiling and downstream spatial analyses. The cohort included 56 post-CCRT ESCC patients, comprising TRG 0 (n = 23), TRG 1 (n = 12), TRG 2 (n = 17), and TRG 3 (n = 4), with the primary comparison defined as TRG 0–1 (n = 35) versus TRG 2–3 (n = 21). (B) Mean cell-type category fractions by TRG group, based on CellDART deconvolution of Visium spots. Fine-grained cell states were collapsed into broad cell categories for visualization. (C) Per-patient log2(effector/TAM) ratio by TRG group; P value by Mann–Whitney U test. (D) Representative Visium maps from a good responder (TRG 0, top row) and a poor responder (TRG 3, bottom row), showing H&E, CancerFinder-defined malignant-cell-enriched annotation, Macro-CXCL5, Tem-GZMK, and EC-FLT1. Scale bars, 500 µm.

CellDART deconvolution showed that TRG 2–3 cores were characterized at the tissue level by a shift toward macrophage-rich, effector-poor, and endothelial-depleted composition (**Figure 1B**). Compared with TRG 0–1 tumors, TRG 2–3 tumors showed lower Tem-GZMK, Temra-CD8, and endothelial fractions together with higher malignant and Macro-CXCL5 fractions and a reduced effector/TAM ratio (**Figure 1C; Supplementary Figure S4**). Representative Visium maps illustrated a good responder with sparse malignant-cell-enriched regions, low Macro-CXCL5 signal, and preserved EC-FLT1 signal, contrasted with a poor responder showing broad malignant-cell-enriched regions, increased Macro-CXCL5, and reduced EC-FLT1 signal (**Figure 1D**).

### Spatial immune landscape remodeling in CCRT-resistant residual tumors: effector exclusion, macrophage accumulation, and microvascular rarefaction

To determine whether TRG-associated immune differences reflected altered localization rather than abundance alone, we calculated the signed distance of each Visium spot to the nearest boundary of the CancerFinder-defined malignant-cell-enriched region. Negative values denoted spots within malignant-cell-enriched regions, whereas positive values denoted surrounding nonmalignant tissue within the same sampled TMA core. Representative signed-distance maps are shown in **Supplementary Figure S5**. This approach was used because local immune exclusion cannot be inferred from abundance alone, particularly in 2-mm TMA cores that sample only a portion of the residual lesion.

Effector immune programs were depleted within malignant-cell-enriched regions of TRG 2–3 tumors. At the tissue level, Tem-GZMK cytotoxic T-cell, Temra-CD8 terminal effector, and DC-CLEC9A cross-presenting dendritic-cell programs were reduced in TRG 2–3 cores, whereas NK-XCL2 showed a similar directional trend (**Figure 2A**). Distance-resolved analyses demonstrated that these effector populations preferentially localized outside malignant-cell-enriched regions in both groups, but that intraregional depletion was more pronounced in TRG 2–3 tumors (**Figure 2B**). Among the 25 cores with residual cancer and analyzable malignant-cell-enriched regions, effector immune programs within malignant-cell-enriched regions were significantly reduced in TRG 2–3 tumors (**Figure 2C**). STopover analysis likewise indicated reduced spatial co-localization between effector immune programs and malignant regions in TRG 2–3 tumors (**Figure 2D**).

**Figure 2.**
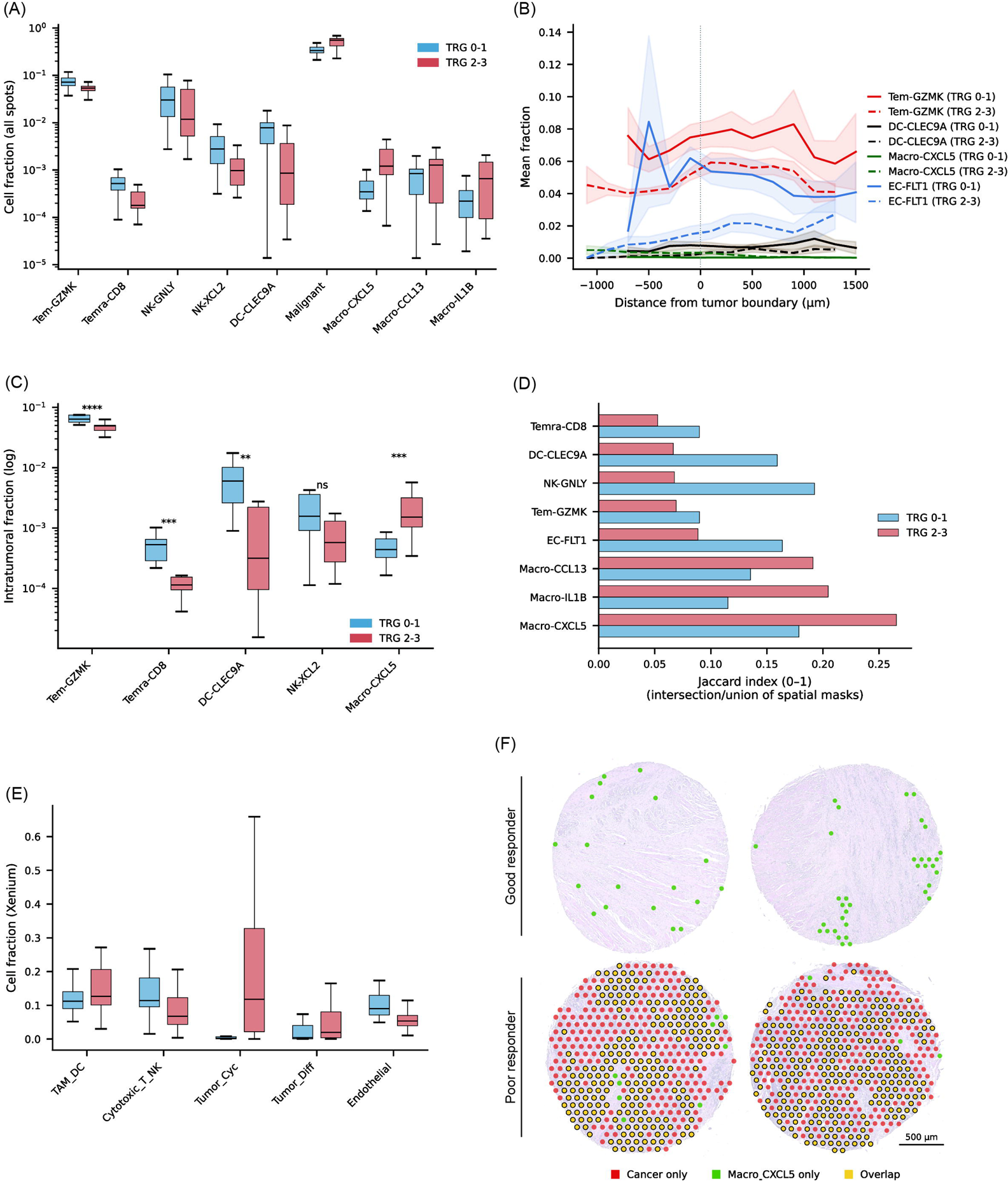
Spatial immune exclusion, macrophage accumulation, and endothelial depletion in chemoradiotherapy-resistant residual ESCC. (A) Patient-level tissue-wide fractions across all Visium spots for selected effector immune, malignant, and macrophage cell states, comparing TRG 0–1 and TRG 2–3 tumors. (B) Distance-resolved mean cell-state fractions relative to the boundary of the CancerFinder-defined malignant-cell-enriched region. Negative distances indicate spots within malignant-cell-enriched regions, whereas positive distances indicate surrounding nonmalignant tissue within the same sampled TMA core. Shaded bands indicate SEM across patients. (C) Patient-level fractions of selected cell states within CancerFinder-defined malignant-cell-enriched regions among residual-cancer cores with analyzable malignant regions (n = 25). Asterisks indicate adjusted significance as shown in the panel. (D) Spatial co-localization between selected cell-state programs and malignant regions, quantified by STopover Jaccard overlap score. Higher values indicate greater spatial overlap with malignant regions. (E) Xenium orthogonal single-cell-resolution refinement showing per-core fractions of major cell populations in serial sections from the same TMA blocks. (F) Representative Visium cores from two good responders (both TRG 0) and two poor responders (TRG 2 and TRG 3) showing CancerFinder-defined malignant spots (Cancer only), Macro-CXCL5-only spots, and overlap between the two. Good responders show minimal overlap, whereas poor responders show broad overlap between malignant and Macro-CXCL5-positive regions. (Scale bars, 500 µm) In box plots, center lines indicate medians and boxes indicate interquartile ranges.

Xenium provided orthogonal single-cell-resolution refinement of this pattern. At the cell level, poor responders showed lower Cytotoxic_T_NK and Endothelial fractions together with higher TAM_DC and tumor-cell fractions, consistent with a macrophage-rich, effector-poor, endothelial-depleted resistant niche (**Figure 2E; Supplementary Figure S6**).

In striking contrast to the spatial exclusion of effector immune cells, macrophage-associated programs were not only preserved but actively enriched in TRG 2–3 tumors. Macro-CXCL5 showed the opposite pattern across the same analytical framework: increased at the tissue level, preferentially concentrated within malignant-cell-enriched regions in distance-resolved analyses, enriched in intratumoral residual disease, and demonstrating strong spatial overlap with malignant regions by STopover (**Figure 2D**). Representative Visium cores visually demonstrated this contrast: good responders (TRG 0) showed minimal Cancer × Macro-CXCL5 overlap with sparse green spots (Macro-CXCL5 only) and limited yellow overlap, whereas poor responders (TRG 2–3) exhibited broad yellow overlap regions indicating extensive macrophage accumulation within malignant-cell-enriched areas (**Figure 2F**). Because the Macro-CXCL5 state in the reference atlas coexpressed SPP1 alongside CXCL5, we interpreted this program as an SPP1⁺/CXCL5⁺ macrophage state within the resistant niche.

A second defining feature of this resistant niche was endothelial depletion consistent with microvascular rarefaction. All four endothelial subtypes identified by CellDART were reduced in TRG 2–3 tumors, accompanied by concordant decreases in multiple pan-endothelial genes (PECAM1, CDH5, CLDN5, VWF, TEK; **Supplementary Figure S7**), while FLT1 gene-level expression was relatively preserved, arguing against selective loss of a single endothelial subset. In the post-CCRT context, such vascular loss is biologically relevant because impaired vascular recovery may aggravate hypoxia, limit drug delivery, and reduce immune-cell trafficking, contributing to persistence of treatment-resistant residual niches. Together, these spatial analyses reveal a coordinated immune-vascular remodeling in CCRT-resistant residual ESCC: effector immune cells are spatially excluded, SPP1⁺/CXCL5⁺ macrophages preferentially accumulate, and the microvasculature is depleted within malignant-cell-enriched regions.

### Single-cell-resolution dissection of CXCL5⁺ and SPP1⁺ macrophage states reveals shared SPP1 expression and immunosuppressive signaling to CD8⁺ T cell

To mechanistically interpret the SPP1⁺/CXCL5⁺ macrophage signal observed in the spatial data, we re-analyzed the published ESCC single-cell RNA-seq atlas (GSE221561) (15), with focus on the myeloid compartment. Unsupervised clustering recovered a coherent monocyte/macrophage population (**Figure 3A**). Subclustering of the monocyte/macrophage compartment resolved a CXCL5⁺ macrophage state and three SPP1⁺ macrophage clusters; remaining clusters were grouped as Monocyte_other. CXCL5⁺ macrophages co-expressed SPP1 alongside CXCL5, whereas SPP1⁺ macrophages expressed SPP1 in the absence of substantial CXCL5 expression (**Figure 3B**). This indicates that CXCL5⁺ macrophages constitute a transcriptional subset of the broader SPP1⁺ macrophage population, providing a single-cell-resolution interpretation of the Macro-CXCL5 program enriched in TRG 2–3 spatial cores.

**Figure 3.**
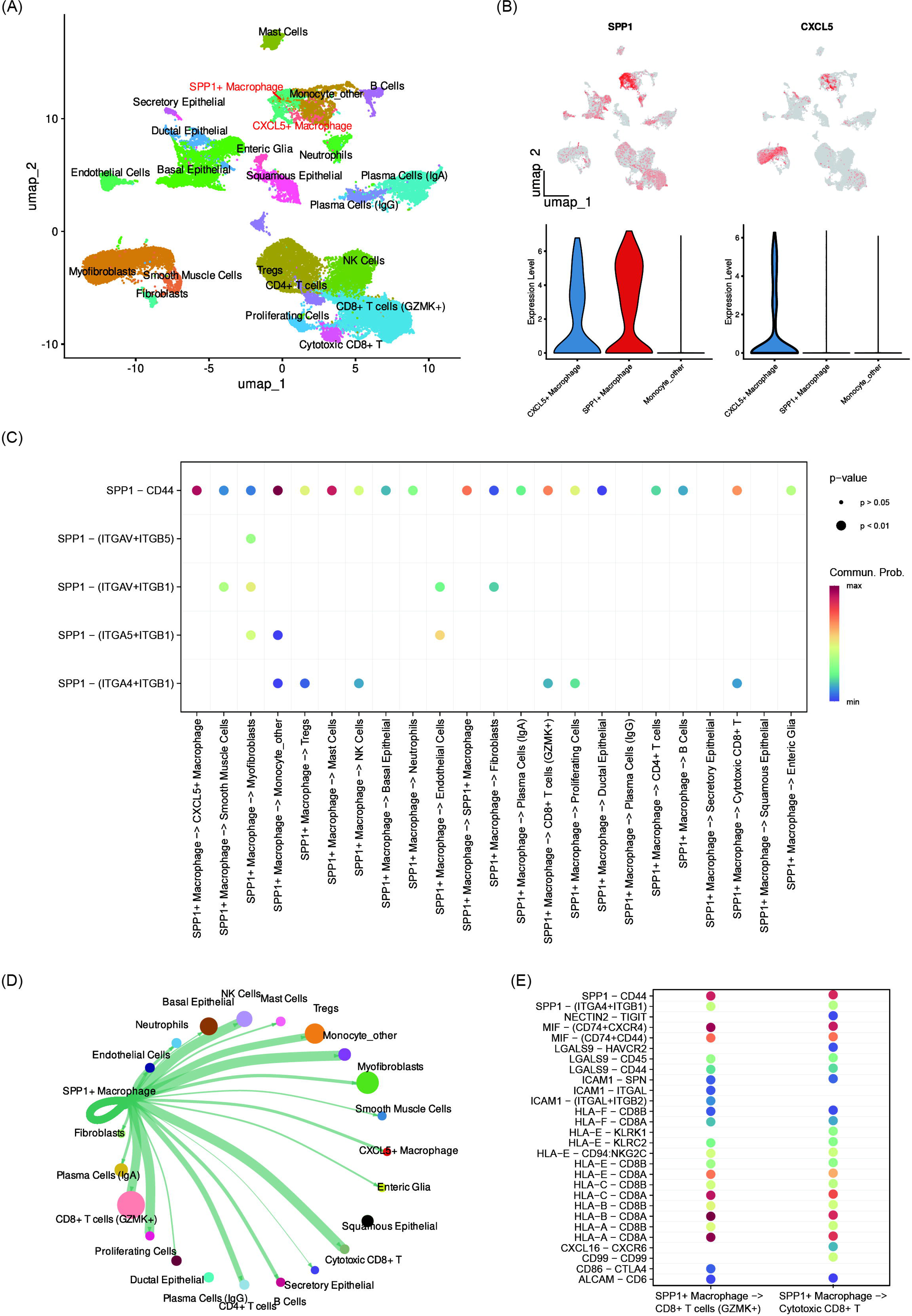
Single-cell-resolution dissection of CXCL5⁺ and SPP1⁺ macrophage states reveals shared SPP1 expression and immunosuppressive signaling to CD8⁺ T cells. (A) UMAP embedding of published ESCC scRNA-seq atlas. (B) SPP1 and CXCL5 expression in macrophage subclusters. (C) CellChat-based SPP1 pathway ligand–receptor interactions from SPP1⁺ macrophages. Dot size represents communication probability; color represents expression level or significance. (D) SPP1 pathway signaling enriched toward stromal and epithelial targets. Line thickness represents communication strength. (E) Immunosuppressive checkpoint signaling from SPP1⁺ macrophages to CD8⁺ T cells. Bubble plots showing ligand–receptor pairs.

CellChat-based ligand–receptor analysis revealed that SPP1⁺ macrophages dominantly signaled through the SPP1 pathway, with broad SPP1–CD44 communication directed at most receiver cell types and additional SPP1–(ITGAV+ITGB5), SPP1–(ITGAV+ITGB1), SPP1–(ITGA5+ITGB1), and SPP1–(ITGA4+ITGB1) interactions (**Figure 3C**) enriched toward stromal and epithelial targets, including myofibroblasts, fibroblasts, and squamous epithelial cells (**Figure 3D**). Of particular relevance to the immune exclusion phenotype, signaling from SPP1⁺ macrophages to CD8⁺ T-cell populations included not only canonical HLA class I–mediated antigen presentation but also a coordinated panel of immunosuppressive checkpoint axes — NECTIN2–TIGIT, CD86–CTLA4, and LGALS9–HAVCR2 (TIM-3) (**Figure 3E**). Collectively, these results provide a mechanistic basis for the spatial co-occurrence of SPP1⁺/CXCL5⁺ macrophage accumulation and effector immune-cell exclusion in CCRT-resistant residual ESCC.

### CCRT-resistant residual tumors show coordinated metabolic and proliferative pathway reprogramming

To determine whether the TRG 2–3 resistant niche was associated with broader biologic reprogramming, we inferred pathway activities from Visium whole-transcriptome data. TRG 2–3 tumors showed coordinated increases in TNFα, VEGF, JAK–STAT, and MAPK activity together with relative reduction of TRAIL signaling (**Figure 4A and Supplementary Figure S8**). HALLMARK analyses identified a parallel proliferative-metabolic program characterized by enrichment of MYC targets, mTORC1 signaling, a second MYC-target program, the p53 pathway, glycolysis, and E2F/G2M-related cell-cycle programs (**Figure 4B and Supplementary Figure S8**).

**Figure 4.**
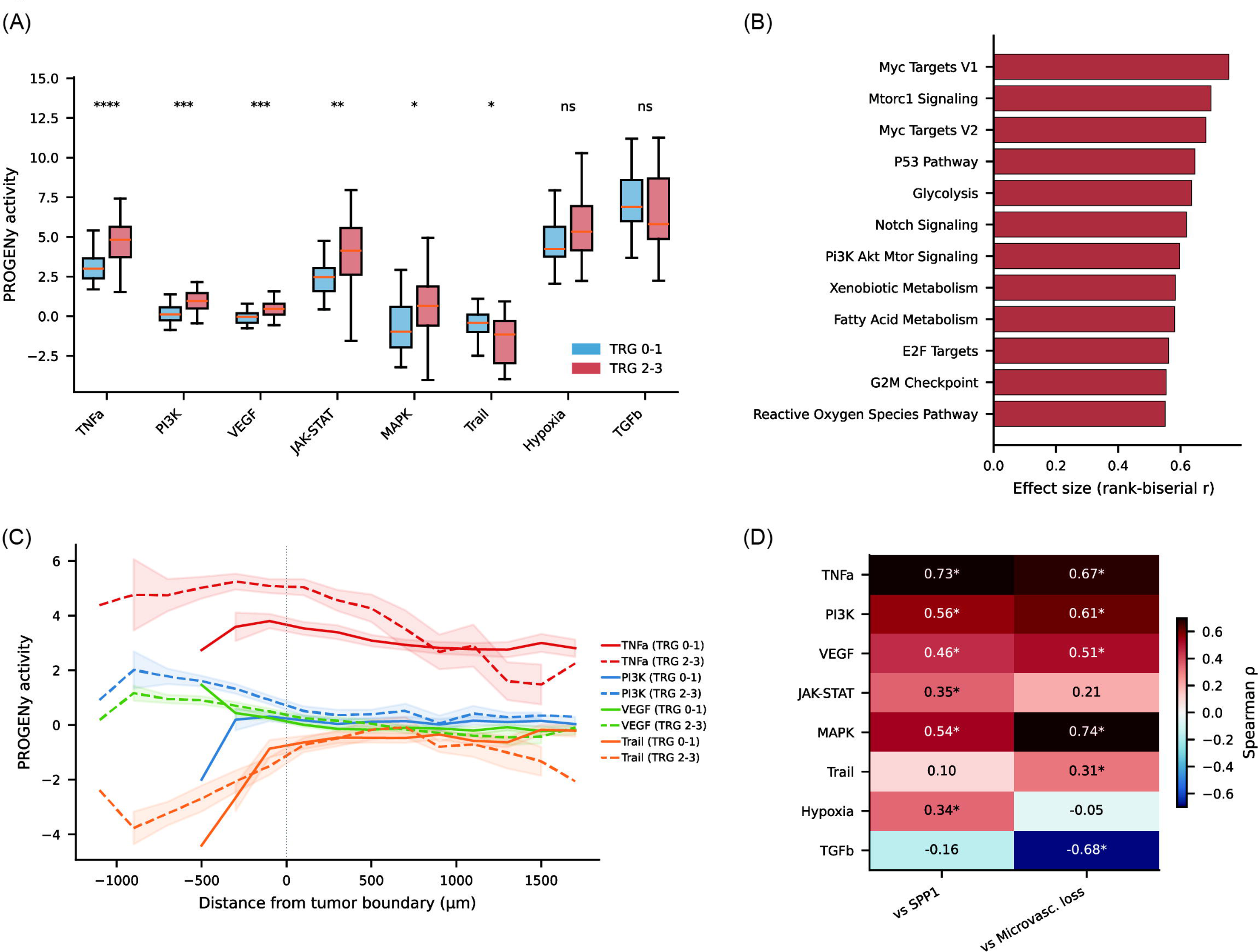
CCRT-resistant residual tumors show coordinated inflammatory, survival, metabolic, and proliferative pathway reprogramming. (A) Patient-level PROGENy pathway activities inferred from Visium whole-transcriptome data, comparing good responders (TRG 0–1; n = 35) and poor responders (TRG 2–3; n = 21). Box plots show medians and interquartile ranges; asterisks indicate significance as shown in the panel. (B) Top differential HALLMARK pathways ranked by rank-biserial effect size. Positive values indicate relative enrichment in TRG 2–3 tumors. (C) Spatial gradients of selected PROGENy pathway activities (TNFα, PI3K, VEGF, and TRAIL) relative to the boundary of CancerFinder-defined malignant-cell-enriched regions. Negative distances indicate spots within malignant-cell-enriched regions, whereas positive distances indicate surrounding nonmalignant tissue within the same sampled TMA core. Line styles denote TRG groups as labeled in the panel, and shaded bands indicate SEM across patients. (D) Patient-level Spearman correlations between pathway activities and resistant-niche features, shown for SPP1 expression and microvascular loss. Positive values indicate higher pathway activity with increasing feature values, whereas negative values indicate inverse correlations. Asterisks indicate significant correlations as shown in the panel.

These pathway differences were not spatially uniform. Several pathway activities showed gradients around local malignant-cell-enriched boundaries, and representative pathway maps illustrated stronger tumor-proximal activity in poor responders than in good responders (**Figure 4C and Supplementary Figure S9**). At the patient level, selected pathway scores correlated positively with SPP1 expression and/or endothelial-loss features, linking inflammatory, survival, and proliferative reprogramming to the macrophage-rich resistant niche (**Figure 4D**). Collectively, these data indicate that CCRT-resistant residual tumors are accompanied by coordinated inflammatory, survival, metabolic, and proliferative reprogramming centered on the SPP1-rich, vascular depleted niche.

### Spatial SPP1-associated ligand–receptor programs link macrophages to tumor cells and CAFs in CCRT-resistant niches

Building on the single-cell prediction that SPP1-associated signaling preferentially targets tumor and stromal receivers, we asked whether the same axes were detectable in space and which cell types expressed candidate SPP1 receptors at baseline. Of the nine Xenium-annotated cell types shown in **Figure 5A**, CAF and tumor epithelial cells (Tumor_Diff, Tumor_Cyc) displayed the highest baseline ITGB1 expression, motivating their selection as plausible SPP1 receiver candidates. Endothelial cells and vSMCs were retained as comparators in the downstream distance-resolved analysis.

**Figure 5.**
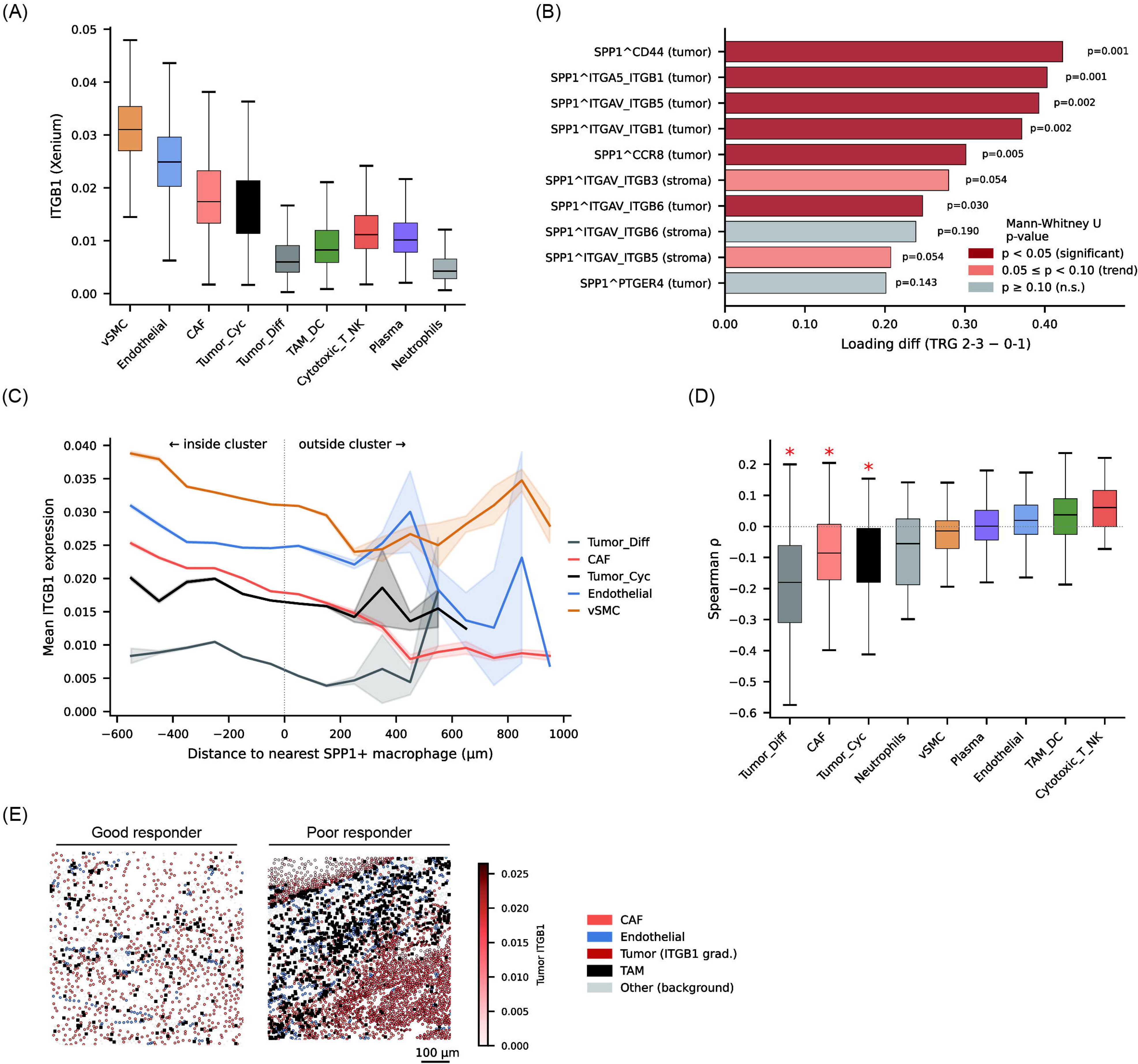
SPP1-associated ligand–receptor programs and candidate receiver-cell biology in CCRT-resistant residual ESCC. (A) Xenium per-cell ITGB1 expression across annotated cell types. Box plots show medians and interquartile ranges. (B) Top-ranked SPP1-containing ligand–receptor pairs identified from Visium-based LIANA analysis, ordered by loading increase in TRG 2–3 versus TRG 0–1 tumors. Labels indicate tumor or stromal spatial domains. P values and significance categories are shown in the panel. (C) Mean Xenium ITGB1 expression as a function of signed proximity to SPP1-high macrophage clusters, shown separately for candidate receiver cell types. (D) Per-patient Spearman correlations between ITGB1 expression and signed proximity to SPP1-high macrophage clusters, stratified by cell type. More negative values indicate higher ITGB1 expression in cells closer to SPP1-high macrophages. (E) Representative Xenium close-up maps from a good responder and a poor responder, illustrating the spatial relationship among TAMs, tumor-cell ITGB1 signal, CAFs, and endothelial cells. These images are shown as single-cell-resolution spatial refinement of candidate receiver-cell biology rather than as direct evidence of causal interaction. Scale bars, 100 µm.

In Visium-based LIANA analyses, SPP1-containing interactions were enriched among tumor-domain ligand–receptor programs in TRG 2–3 tumors. Among the top-ranked SPP1-containing pairs, SPP1–CD44, SPP1–ITGA5/ITGB1, SPP1–ITGAV/ITGB5, and SPP1– ITGAV/ITGB1 were the most enriched tumor-domain interactions, whereas stromal-domain SPP1 interactions were weaker but detectable (**Figure 5B**). The broader ligand–receptor landscape is shown in **Supplementary Figure S10**, and the top-ranked SPP1-containing and ITGB1-containing ligand–receptor pairs are listed in Supplementary Tables S2 and S3, respectively.

We then used Xenium to refine candidate receiver cell types at single-cell resolution. For each Xenium cell, we calculated the signed distance to the nearest SPP1-high macrophage cluster boundary and examined ITGB1 expression as a function of this signed proximity, separately by cell type. Tumor_Diff and CAF populations showed higher ITGB1 expression in cells located closer to SPP1-high macrophages, whereas endothelial cells did not show a comparable gradient (**Figure 5C**).

To quantify this pattern at the patient level, we calculated within each patient and cell type the Spearman correlation between ITGB1 expression and signed distance to the nearest SPP1-high macrophage cluster boundary. Negative correlations indicate higher ITGB1 expression in cells closer to SPP1-high macrophages. Tumor_Diff, CAF, and Tumor_Cyc compartments showed the strongest negative correlation distributions, whereas Endothelial, TAM_DC, and Cytotoxic_T_NK populations did not (**Figure 5D**). Representative Xenium close-up maps further illustrated dense TAM accumulation adjacent to ITGB1-high tumor cells in a poor responder, compared with sparse TAM infiltration and low tumor ITGB1 signal in a good responder (**Figure 5E**). Together with the single-cell ligand–receptor inference, these data support a model in which SPP1-associated macrophage programs are preferentially linked to tumor and fibroblast receiver compartments within resistant residual ESCC.

### Resistant-niche features show supportive associations with disease-free survival in the SNUH cohort, with orthogonal supportive context from TCGA ESCC

Having identified SPP1-associated macrophage accumulation and microvascular rarefaction as the two major biologic axes of the TRG 2–3 resistant niche, we next asked whether these emergent features were associated with clinical outcome. Within the SNUH cohort, core-level SPP1 expression increased across TRG groups and was higher in TRG 2–3 tumors (**Figure 6A**). In univariable Cox analysis, higher SPP1 expression was associated with shorter DFS (HR per 1-SD increase, 1.65; 95% CI, 1.15–2.38; **Figure 6E**), and Kaplan–Meier analysis showed significantly worse DFS in the high-SPP1 group compared with the low-SPP1 group by median split (log-rank P = 0.0213; **Figure 6B**). Lower endothelial abundance showed an even stronger association with shorter DFS (HR, 2.04; 95% CI, 1.36–3.07; Figure 6E), with the microvascular-loss score conferring markedly worse DFS by median split (log-rank P = 0.0004; **Figure 6C**). A combined SPP1 + microvascular-loss score further stratified DFS (log-rank P = 0.0002; **Figure 6D**). In a joint multivariable model including SPP1, microvascular loss, and ypStage, microvascular loss and ypStage retained significant associations whereas SPP1 did not, suggesting that these variables capture overlapping aspects of the same unfavorable residual-disease biology (**Figure 6F**). Sensitivity analyses across alternative TRG groupings and subgroups are shown in **Supplementary Figure S11**. Similar directional patterns were observed for LRFS, DMFS, and OS, with results summarized in **Supplementary Figures S12–S14**.

**Figure 6.**
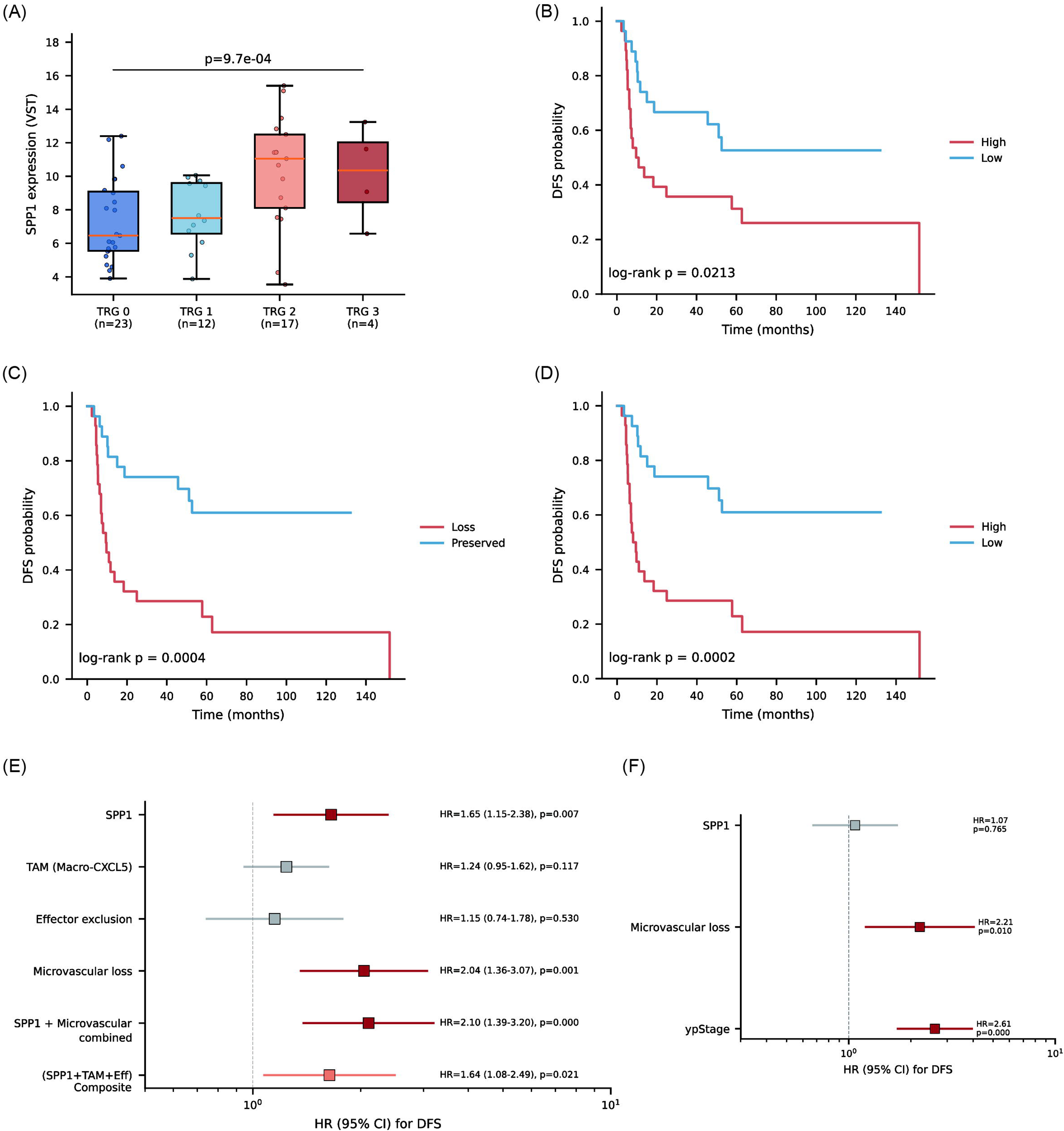
Supportive clinical relevance of resistant-niche features in post-CCRT ESCC. (A) Core-level SPP1 expression by tumor regression grade (TRG) group in the SNUH cohort. (B) Kaplan–Meier curves for disease-free survival (DFS) stratified by median-split SPP1 expression. Median split was used for descriptive visualization. (C) Kaplan–Meier curves for DFS stratified by the median-split microvascular-loss score. Median split was used for descriptive visualization. (D) Kaplan–Meier curves for DFS stratified by the combined SPP1 + microvascular-loss score. Median split was used for descriptive visualization. (E) Univariable Cox proportional hazards models for DFS using continuous z-scored resistant-niche features, including SPP1, TAM (Macro-CXCL5), effector exclusion, microvascular loss, the combined SPP1 + microvascular-loss score, and a broader composite score. Hazard ratios are shown with 95% confidence intervals. (F) Multivariable Cox proportional hazards model for DFS including SPP1, microvascular loss, and ypStage. Hazard ratios are shown with 95% confidence intervals; continuous variables were modeled per 1-standard-deviation increase.

To externally contextualize the SPP1 association, we analyzed the TCGA ESCC cohort (TCGA-ESCA PanCancer Atlas 2018, squamous histology; n = 63 evaluable patients). In this external orthogonal cohort, higher SPP1 expression showed a trend toward shorter DFS by median split (log-rank P = 0.089; **Supplementary Figure S15A**), directionally consistent with the SNUH cohort. Although the association did not reach statistical significance in this smaller cohort, the directional consistency provides supportive external context for the biologic relevance of SPP1 in ESCC. No significant SPP1–OS association was observed in TCGA (**Supplementary Figure S15B**). Because TCGA-ESCA is not enriched for post-CCRT residual disease, these findings should be interpreted as orthogonal context rather than direct validation of the post-CCRT residual state.

Overall, these data support the clinical relevance of the spatially defined resistant niche in the SNUH cohort, while the TCGA analysis provides orthogonal supportive context for the broader biologic relevance of SPP1 in ESCC. These findings should still be interpreted as hypothesis-generating with respect to formal biomarker development.

## DISCUSSION

We used paired Visium and Xenium spatial transcriptomics, complemented by single-cell RNA-seq re-analysis and external TCGA cohort survival analysis, to define the tumor microenvironmental architecture of post-CCRT residual ESCC, focusing on TRG 2–3 tumors as a model of chemoradiotherapy-resistant disease. Rather than a change in any single compartment, resistant residual tumors were characterized by a coordinated spatial niche comprising effector immune-cell exclusion from malignant-cell-enriched regions, accumulation of SPP1⁺/CXCL5⁺ macrophage programs, endothelial depletion consistent with microvascular rarefaction, and proliferative-metabolic pathway reprogramming. Single-cell-resolution analyses showed that CXCL5⁺ macrophages constitute a transcriptional subset of the broader SPP1⁺ macrophage population, that SPP1-associated signaling is preferentially oriented toward tumor and CAF receivers via CD44- and integrin-related axes, and that SPP1⁺ macrophages additionally engage immune-checkpoint signaling toward CD8 T cells. Higher SPP1 expression was associated with shorter DFS both in the SNUH cohort and in the independent TCGA ESCC cohort.

An important aspect of our findings is that poor-response biology was expressed spatially. Effector immune programs were not simply reduced in bulk but were preferentially excluded from malignant-cell-enriched regions. This distinction matters because abundance-only measurements can underestimate biologically meaningful immune exclusion. In residual disease after CCRT, where local tumor persistence and recurrence may depend on the survival of small malignant-cell-rich niches, the spatial positioning of cytotoxic T cells, NK cells, and cDC1-like cells is likely more informative than tissue-wide abundance alone (4, 9, 27). Importantly, the single-cell–resolution identification of NECTIN2–TIGIT, CD86–CTLA4, and LGALS9–HAVCR2 as prominent SPP1⁺ macrophage → CD8 T-cell axes provides a candidate mechanistic explanation for this exclusion phenotype: CD8 T cells that do reach the macrophage-dense, SPP1-rich niche are likely to encounter a coordinated immunosuppressive checkpoint milieu. This observation also highlights potential rationales for combining checkpoint inhibition with macrophage-targeted strategies in the post-CCRT residual setting.

SPP1⁺ tumor-associated macrophages have emerged across multiple cancers as immunosuppressive and tumor-supportive populations, often linked to fibroblast-rich and immune-excluded tissue states (24–26). In the present study, however, SPP1 was not a prespecified starting hypothesis; rather, it emerged from the spatial comparison of good and poor responders, was confirmed at single-cell resolution as the dominant signaling backbone of the CXCL5⁺ and SPP1⁺ macrophage states and was independently validated as an unfavorable DFS marker in the TCGA ESCC cohort. The convergence of spatial, single-cell, and external survival data points to a macrophage-centered, SPP1-anchored interaction program that is both spatially organized and clinically informative in residual ESCC. The relative attenuation of SPP1 in our SNUH joint multivariable model is consistent with the interpretation that SPP1, microvascular loss, and ypStage capture overlapping facets of the same unfavorable residual-disease biology rather than statistical independence.

A second major biologic axis in our data was microvascular rarefaction. This finding is important not simply because endothelial abundance may carry prognostic information, but because vascular loss itself may help shape the resistant residual niche after chemoradiotherapy. Radiation-induced endothelial injury and impaired vascular recovery can worsen hypoxia, restrict drug delivery, and reduce immune-cell trafficking, thereby reinforcing survival of unfavorable post-treatment niches (10, 28–31). The broad reduction we observed across endothelial subtypes and pan-endothelial markers is more consistent with global microvascular attrition than with selective remodeling of one endothelial subset.

From a translational perspective, these findings provide a spatially resolved framework for understanding why some ESCCs persist after neoadjuvant CCRT. The convergence of macrophage-centered SPP1-associated interactions, immune-checkpoint-mediated CD8 T-cell suppression, and impaired vascular recovery defines a biologically coherent set of candidate axes for residual-disease persistence. These observations may help prioritize future mechanistic studies and post-CCRT combination-treatment hypotheses, including SPP1- or CD44-axis-directed strategies, macrophage repolarization, vascular normalization, and immune-checkpoint blockade in the residual disease setting. Any future clinical application of SPP1- or vascular-marker evaluation, including immunohistochemistry-based readouts, will require independent validation and prospective study design.

This study has several limitations. First, it is a single-center retrospective discovery cohort. Second, the use of one 2-mm TMA core per patient allows controlled patient-level comparison but does not capture the full architectural heterogeneity of the residual tumor bed. Third, matched pre-CCRT tissue was not available, so baseline versus treatment-induced features could not be distinguished. Fourth, our findings are based on spatial association and ligand–receptor inference rather than direct functional perturbation and therefore require additional functional validation. Fifth, Xenium was generated from serial sections of the same TMA blocks and was used as an orthogonal high-resolution refinement layer rather than as a fully independent validation cohort. Sixth, the single-cell re-analysis used the same atlas (GSE221561) that served as the spatial deconvolution reference and is therefore best interpreted as single-cell-resolution mechanistic dissection rather than as independent dataset validation. Finally, the TCGA ESCC cohort is not enriched for post-CCRT residual disease, and the SPP1 sensitivity-scanned cutoff is reported as supportive evidence; both points should be borne in mind when interpreting the external survival findings.

In summary, chemoradiotherapy-resistant residual ESCC is characterized by a spatially organized tumor microenvironmental niche involving effector immune exclusion, SPP1-associated macrophage accumulation with intrinsic immunosuppressive checkpoint signaling toward CD8 T cells, microvascular rarefaction, and proliferative-metabolic reprogramming. These findings define a macrophage-centered, vasculature-depleted resistant-niche model and identify macrophage–tumor/stromal interactions, immune-checkpoint engagement, and vascular injury or incomplete recovery as candidate axes for overcoming residual disease after CCRT in ESCC.

## Declarations

### Ethics approval and consent to participate

This study was approved by the Institutional Review Board of Seoul National University Hospital (IRB approval number: [J-2105-156-1221], approval date: [2022-06-20]) and conducted in accordance with the Declaration of Helsinki. The requirement for informed consent from individual patients was waived by the Institutional Review Board.

### Consent for publication

Not applicable

### Financial support

This research was supported by the National Research Foundation of Korea (NRF) grant funded by the Korea government (Ministry of Science and ICT, MSIT) (No. RS-2025-00523325) to B.H.Kim.

### Conflict of interest

K.J.N. is a cofounder and shareholder of Portrai, Inc (Republic of Korea) and received a research grant from Inocras (United States). H.C. is a cofounder and shareholder of Portrai, Inc (Republic of Korea). The remaining authors declare no competing interests.

### Data availability

Processed Visium and Xenium AnnData objects of this cohort and clinical metadata necessary to reproduce the SNUH cohort survival analyses are available to qualified investigators on request, subject to institutional data-sharing policies. The single-cell RNA sequencing data is publicly available through GSE221561. The TCGA-ESCA PanCancer Atlas 2018 dataset is publicly available through cBioPortal.

