## Supplementary Figures for "Spatial tumor microenvironmental architecture of chemoradiotherapy-resistant residual esophageal squamous cell carcinoma"

**Supplementary Tables**

### Supplementary Figure Legends

Supplementary Figure 1

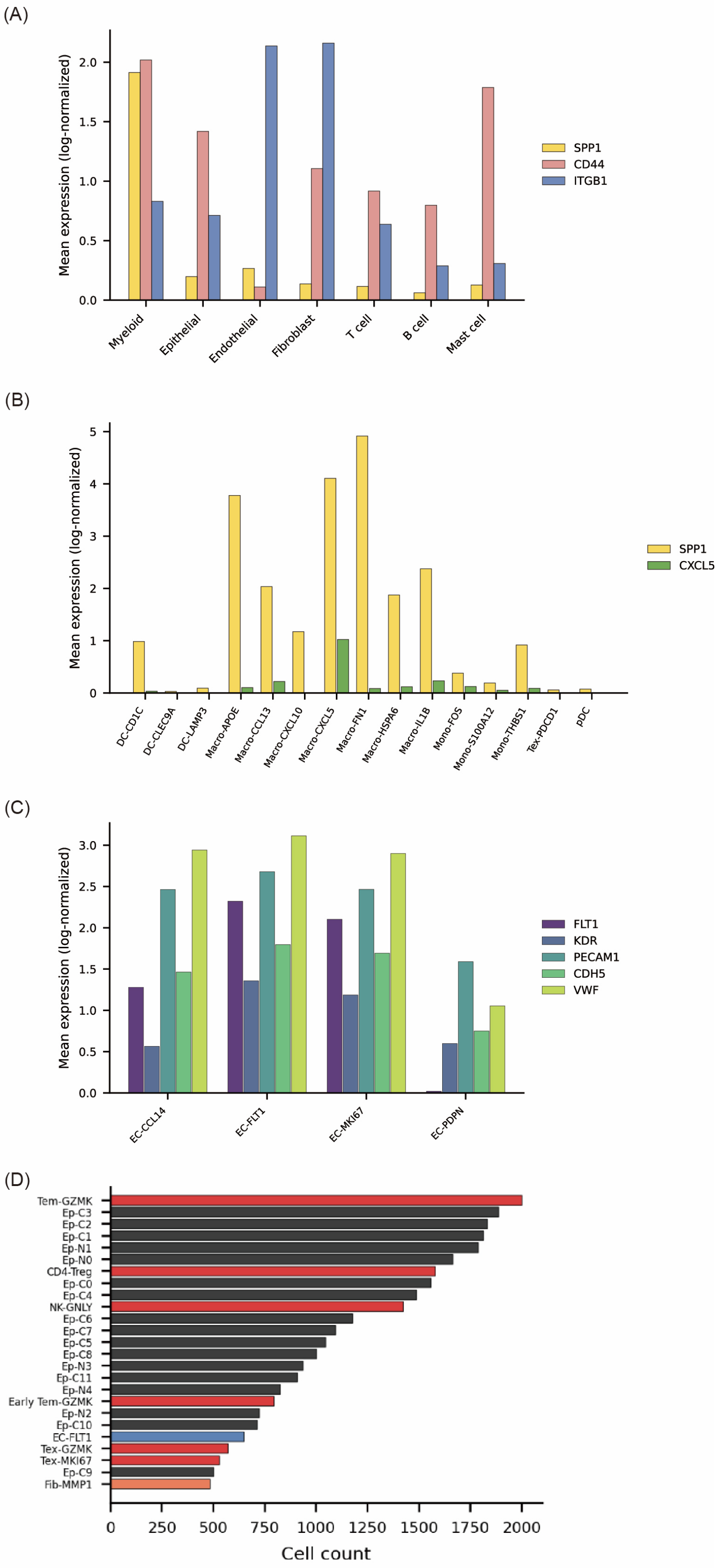

**Supplementary Figure 1. Published ESCC single-cell RNA-seq reference context for SPP1-associated macrophage and endothelial annotations.**

(A) Mean expression (log-normalized) of SPP1, CD44, and ITGB1 across major cell classes in the published ESCC single-cell RNA-seq atlas (GSE221561). SPP1 expression is concentrated in the myeloid compartment, whereas CD44 and ITGB1 are distributed across multiple stromal, epithelial, and immune lineages. (B) Mean expression (log-normalized) of SPP1 and CXCL5 across annotated myeloid subtypes in GSE221561, showing enrichment of both genes in the Macro-CXCL5 state and supporting interpretation of the spatial Macro-CXCL5 program as part of a broader SPP1-associated macrophage continuum. (C) Mean expression (log-normalized) of FLT1, KDR, PECAM1, CDH5, and VWF across endothelial subtypes in GSE221561, providing orthogonal support for endothelial subtype annotation and vascular-marker interpretation in the spatial analyses. (D) Top annotated subtype counts in the published ESCC single-cell RNA-seq atlas, shown as reference context for the cellular composition of GSE221561.

Supplementary Figure 2

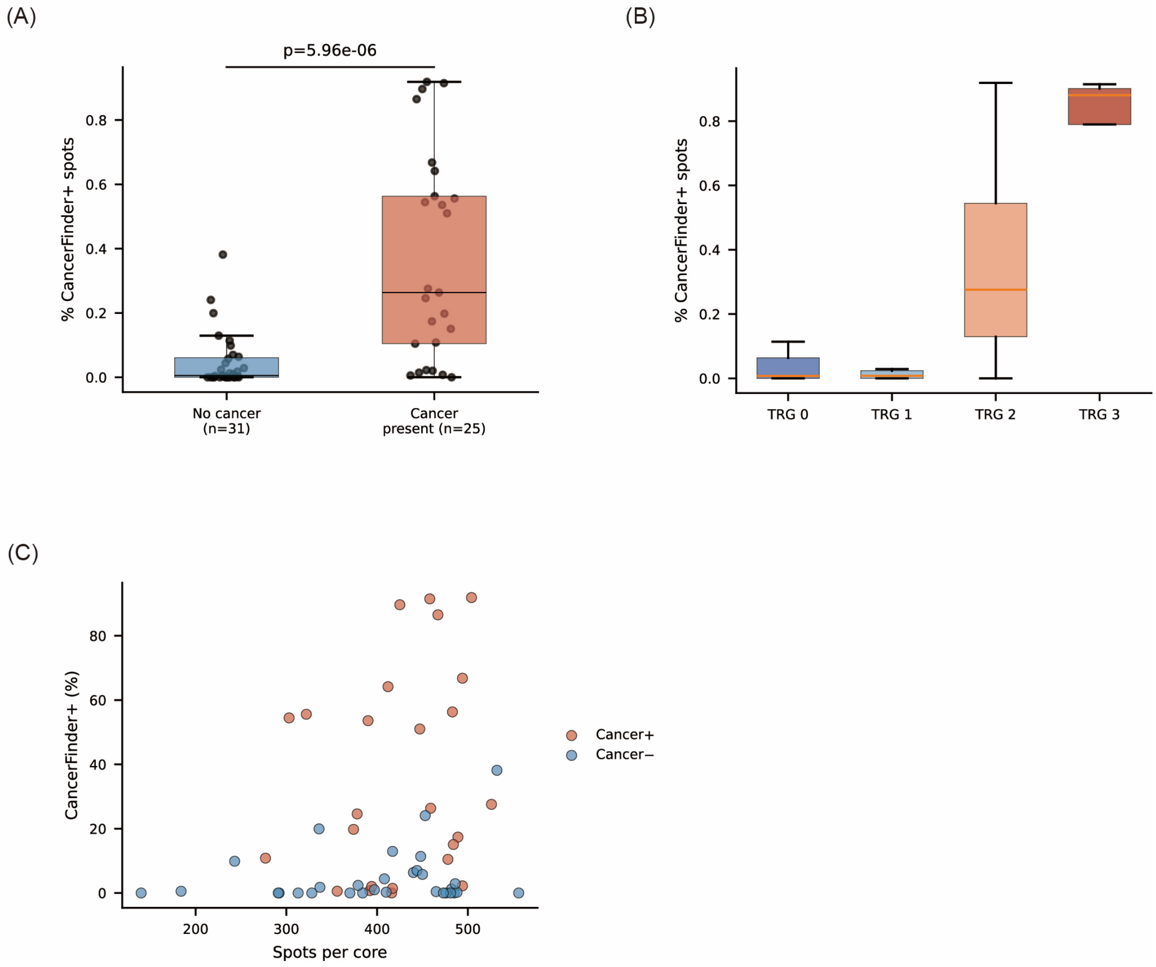


**Supplementary Figure 2. Concordance of CancerFinder-derived malignant-region annotation with pathologist review in post-CCRT ESCC Visium cores.**

(A) Percentage of CancerFinder-positive spots in cores without histologic residual cancer (n = 31) versus cores with pathologist-confirmed residual cancer (n = 25). Each point represents one Visium core; P value by Mann–Whitney U test. (B) Distribution of the CancerFinder-positive spot fraction across tumor regression grade (TRG) 0, 1, 2, and 3 cores. (C) Relationship between total spot number per core and the percentage of CancerFinder-positive spots. Points are colored by pathologist-defined cancer-present status (Cancer = Y vs Cancer = N), allowing visual comparison of CancerFinder-positive fraction across a range of core sizes.

Together, these analyses support the use of CancerFinder to define malignant-cell-enriched regions for downstream spatial analyses. In box plots, center lines indicate medians and boxes indicate interquartile ranges.

Supplementary Figure 3

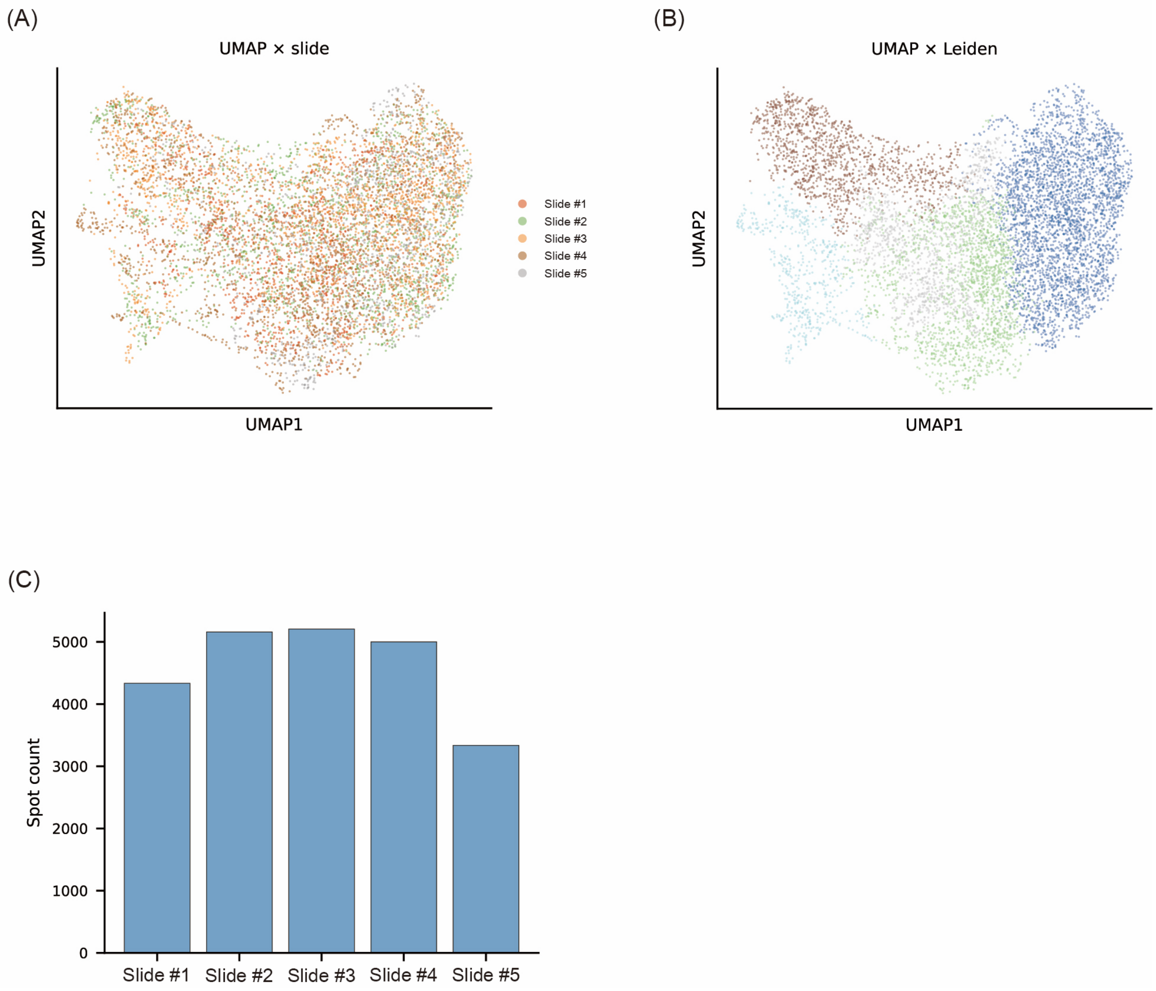


**Supplementary Figure 3. STAligner spatial integration quality control for the multi-slide Visium FFPE dataset.**

(A) UMAP embedding of all Visium spots after STAligner integration, colored by slide identity (Slide #1–Slide #5). Each point represents one Visium spot. (B) The same integrated UMAP embedding colored by STAligner Leiden cluster assignment, illustrating the major integrated spot communities used for downstream analyses. (C) Number of Visium spots contributed by each slide after quality control and integration. Panels A–C provide quality-control context for the integrated multi-slide Visium dataset used in downstream analyses.

Supplementary Figure 4

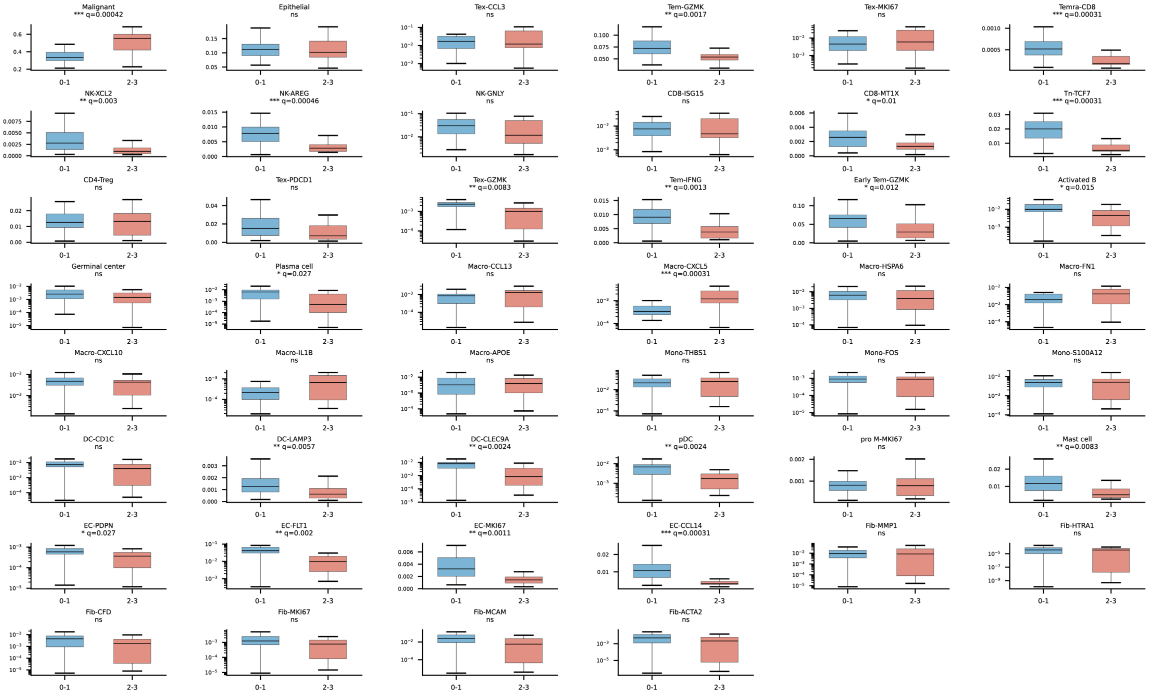


**Supplementary Figure 4. Full patient-level distribution of all 46 CellDART-inferred cell states by TRG group.**

Patient-level mean fractions of all 46 deconvolved cell states across all Visium spots, comparing good responders (TRG 0–1) and poor responders (TRG 2–3). Each box plot represents one cell state. P values were calculated using the Mann–Whitney U test and adjusted by the Benjamini–Hochberg method within the full cell-state family; adjusted q values are shown above individual panels. Box plots show medians and interquartile ranges. This figure provides the full cell-state context underlying the selected cell populations highlighted in the main figures.

Supplementary Figure 5

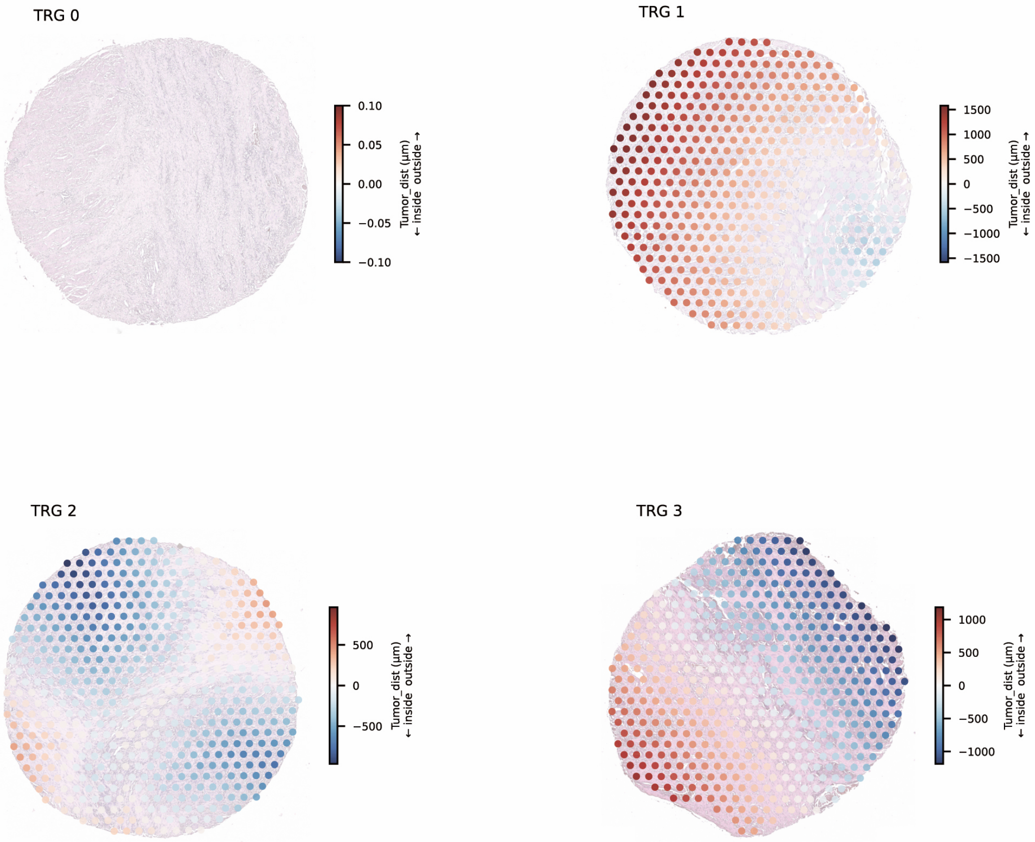


**Supplementary Figure 5. Representative signed-distance maps relative to CancerFinder-defined malignant-cell-enriched regions across TRG groups.**

(A) Representative TRG 0 core. (B) Representative TRG 1 core. (C) Representative TRG 2 core. (D) Representative TRG 3 core. Spots are overlaid on H&E backgrounds and colored by signed distance to the nearest boundary of the CancerFinder-defined malignant-cell-enriched region. Negative values indicate spots within malignant-cell-enriched regions, positive values indicate surrounding nonmalignant tissue within the same sampled TMA core, and zero denotes the local boundary. Distances are shown in micrometers.

Supplementary Figure 6

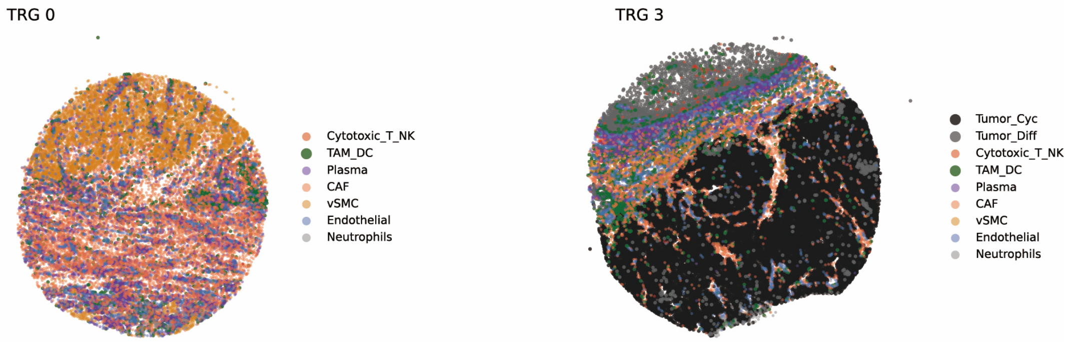


**Supplementary Figure 6. Representative Xenium whole-core cell-type distribution in good and poor responders.**

(A) Representative good responder core (TRG 0). (B) Representative poor responder core (TRG 3). Xenium cells are colored by major cell-type annotation, including Tumor_Cyc, Tumor_Diff, Cytotoxic_T_NK, TAM_DC, Plasma, CAF, vSMC, Endothelial, and Neutrophils. These representative whole-core views provide single-cell-resolution spatial context for the broader compositional differences between good and poor responders.

Supplementary Figure 7

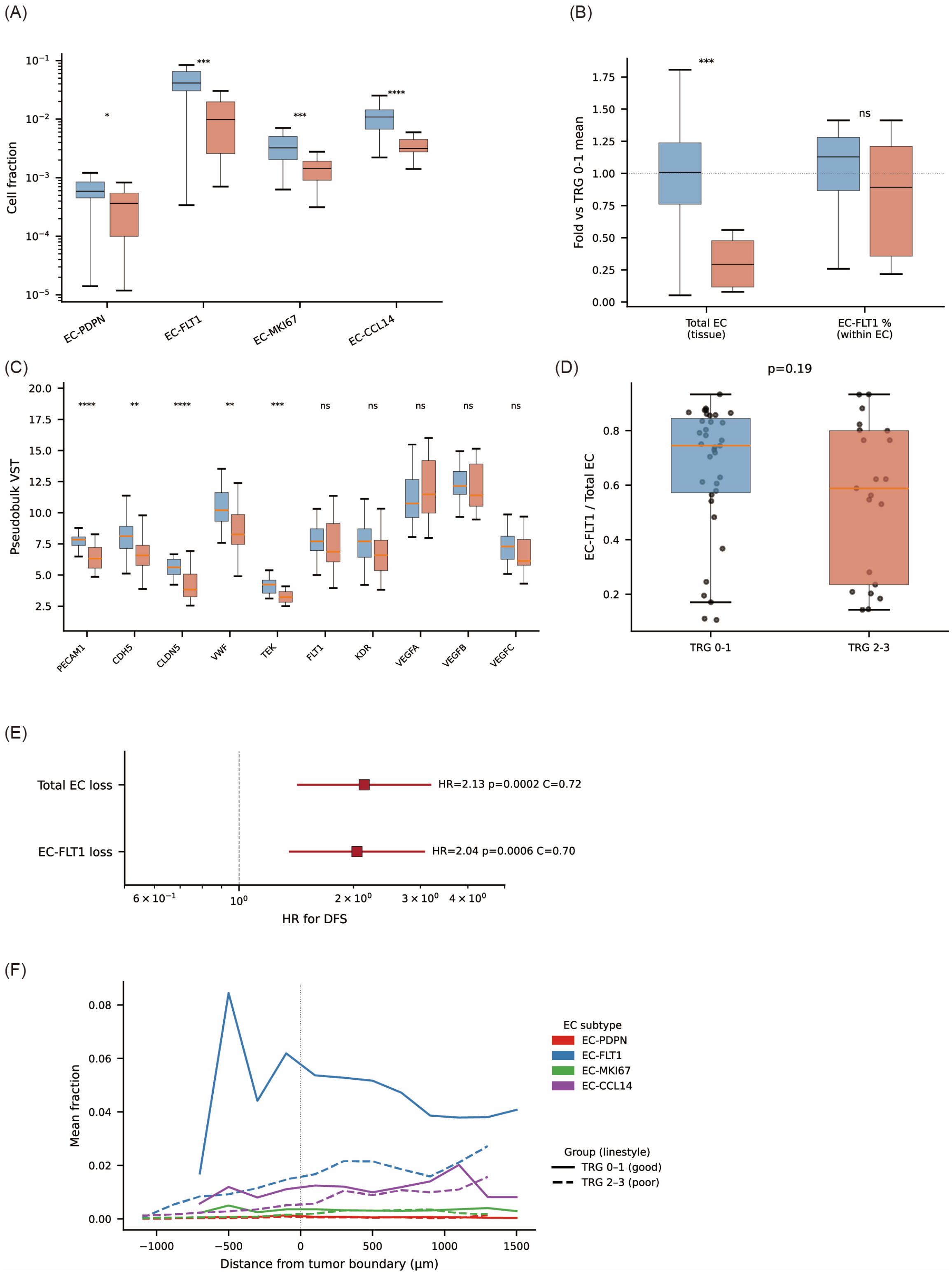

**Supplementary Figure 7. Endothelial depletion and microvascular rarefaction in chemoradiotherapy-resistant residual ESCC.**

(A) Patient-level fractions of the four deconvolved endothelial subtypes (EC-PDPN, EC-FLT1, EC-MKI67, and EC-CCL14) across all Visium spots, comparing TRG 0–1 and TRG 2–3 tumors. (B) Fold change relative to the TRG 0–1 mean for total endothelial abundance and for EC-FLT1 proportion within the endothelial compartment, illustrating overall endothelial loss with relative preservation of EC-FLT1 composition. (C) Patient-level pseudobulk VST-normalized expression of pan-endothelial and VEGF-axis genes. (D) EC-FLT1 fraction normalized to total endothelial abundance, showing preserved within-endothelial EC-FLT1 composition despite overall endothelial depletion. (E) Univariable Cox proportional hazards models for disease-free survival using total endothelial loss and EC-FLT1 loss as continuous predictors; hazard ratios are shown with 95% confidence intervals, with concordance indices annotated in the panel. (F) Distance-resolved mean fractions of the four endothelial subtypes relative to the boundary of CancerFinder-defined malignant-cell-enriched regions. Colors denote endothelial subtypes, and line styles denote TRG groups as labeled in the panel.

These analyses support the interpretation that TRG 2–3 residual tumors exhibit broad endothelial depletion consistent with microvascular rarefaction rather than selective loss of a single endothelial subtype. In box plots, center lines indicate medians and boxes indicate interquartile ranges.

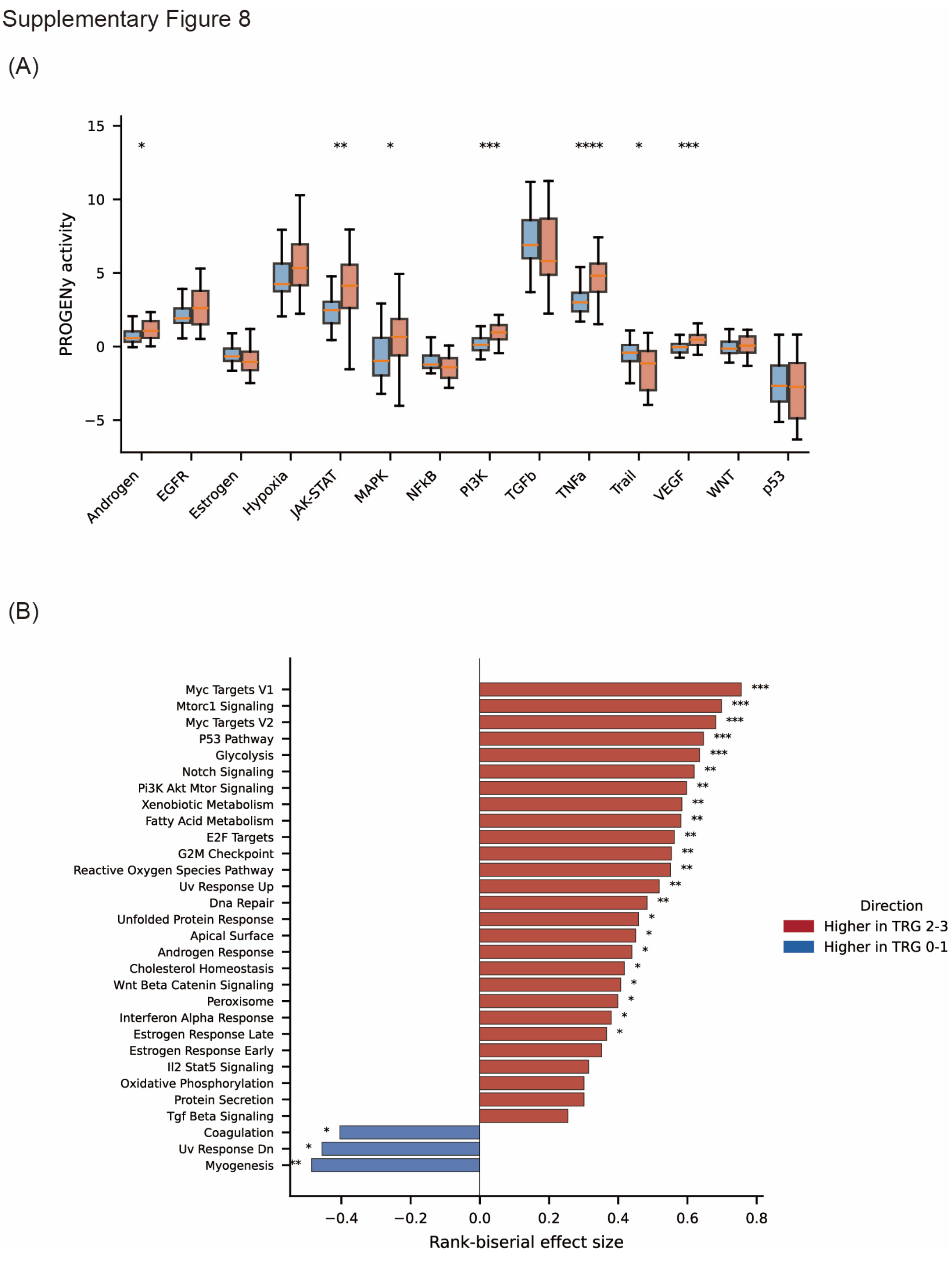


**Supplementary Figure 8. Full PROGENy and top differential HALLMARK pathway results by TRG group.**

(A) Patient-level PROGENy activities for all 14 pathways inferred from Visium whole-transcriptome data, comparing TRG 0–1 and TRG 2–3 tumors. Statistical comparisons used the Mann–Whitney U test with Benjamini–Hochberg correction within the PROGENy pathway family; asterisks indicate adjusted significance as shown in the panel. (B) Top 30 HALLMARK pathways ranked by absolute rank-biserial effect size. Positive values indicate relative enrichment in TRG 2–3 tumors, whereas negative values indicate relative enrichment in TRG 0–1 tumors. Colors denote direction of enrichment as annotated in the panel.

Supplementary Figure 9

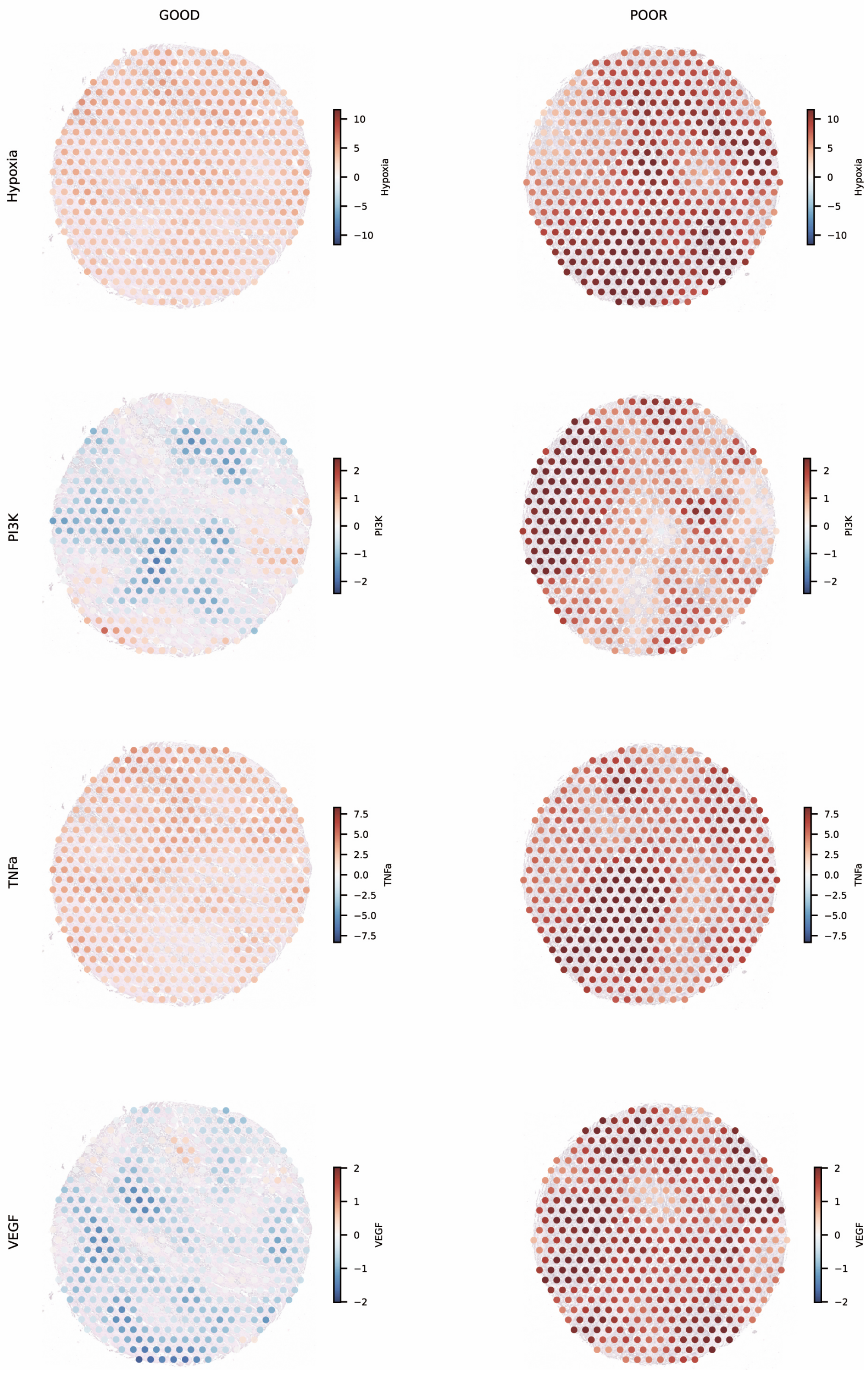

**Supplementary Figure 9. Representative spatial maps of pathway activity in good and poor responders.**

Top row, representative good responder core (TRG 0; P-25-R-U8-B29-54-V_C04). Bottom row, representative poor responder core (TRG 3; P-25-R-U8-B28-54-V_C01). Visium spots are overlaid on H&E backgrounds and colored by inferred PROGENy pathway activity for TNFα, VEGF, Hypoxia, and PI3K. Color bars indicate relative pathway activity as shown in each panel.

These representative maps illustrate broader positive TNFα, Hypoxia, PI3K, and VEGF activity in the poor responder than in the good responder, consistent with the pathway-level differences observed in the patient-level analyses.

Supplementary Figure 10

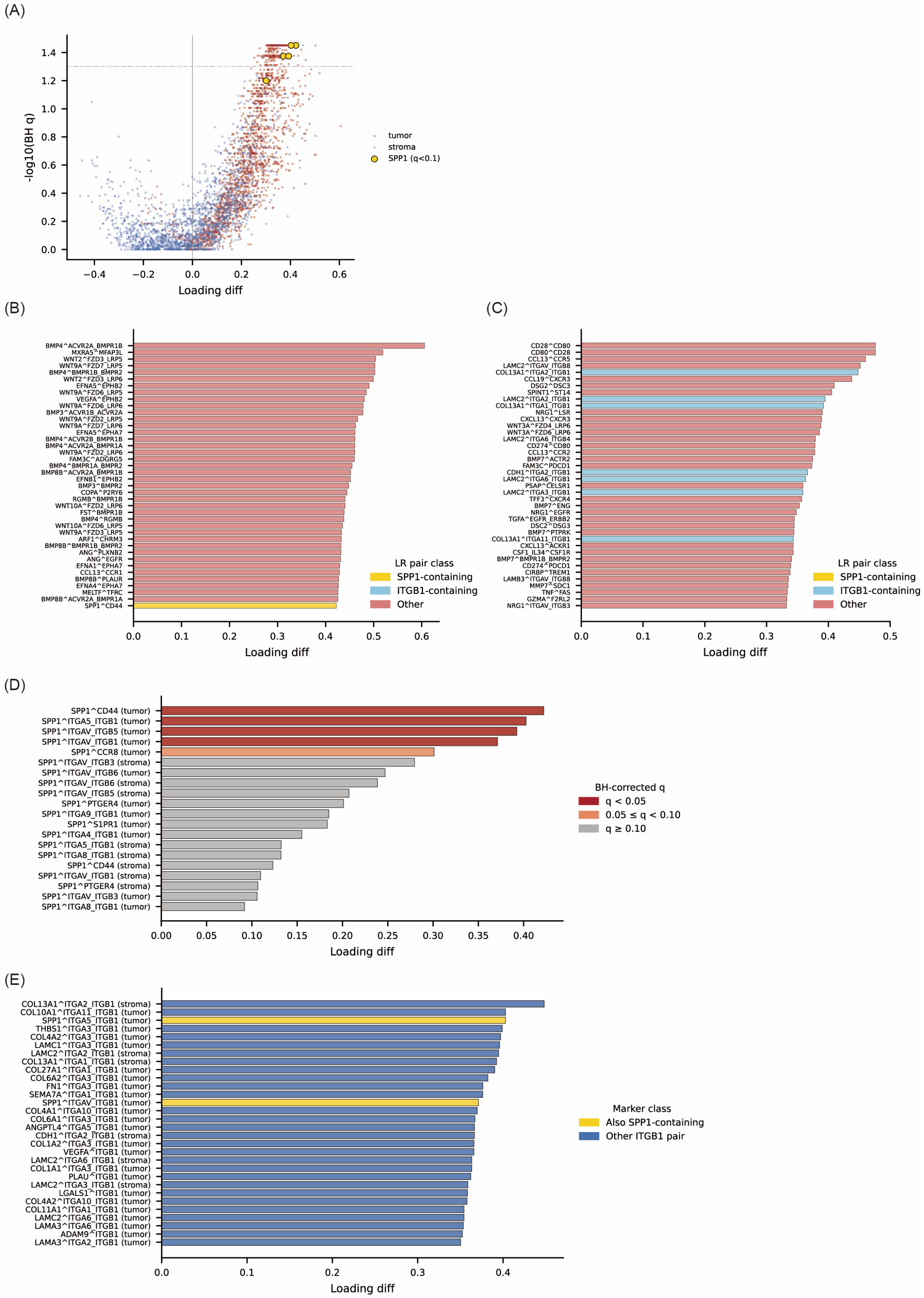

**Supplementary Figure 10. Full LIANA ligand–receptor interaction landscape across tumor and stromal domains.**

(A) Volcano plot of all 3,809 ligand–receptor pairs with nonzero loading, showing loading difference (TRG 2–3 minus TRG 0–1) versus −log10(BH-adjusted q value). Tumor-domain and stroma-domain pairs are indicated separately, and SPP1-containing pairs are highlighted as annotated. (B) Top 40 tumor-domain ligand–receptor pairs ranked by loading increase in TRG 2–3 versus TRG 0–1. Bar colors follow the panel legend, distinguishing SPP1-containing, ITGB1-containing, and other pairs. (C) Top 40 stroma-domain ligand–receptor pairs ranked by loading increase in TRG 2–3 versus TRG 0–1. (D) All SPP1-containing ligand–receptor pairs (n = 20), ranked by loading difference and annotated by spatial domain. (E) Top 30 ITGB1-containing ligand–receptor pairs. SPP1–ITGB1-containing pairs are highlighted as annotated in the panel.

Supplementary Figure 11

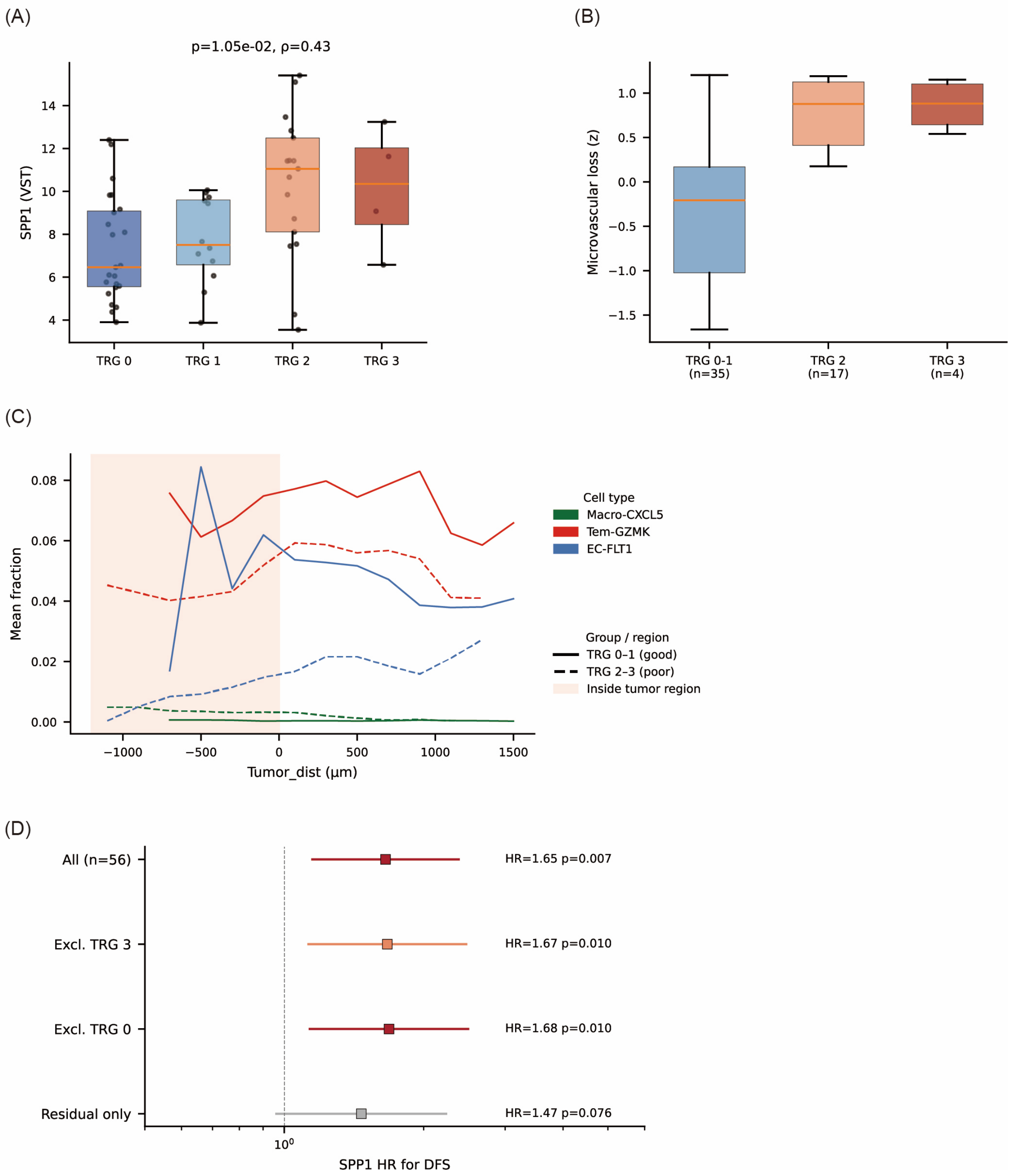


**Supplementary Figure 11. Sensitivity analyses for resistant-niche features and outcome associations.**

(A) SPP1 expression across the four TRG groups (TRG 0, 1, 2, and 3), shown to illustrate the graded increase across pathologic response categories. (B) Microvascular-loss score across TRG 0–1, TRG 2, and TRG 3 groups. (C) Distance-resolved mean fractions of selected features under an alternative analytic setting, showing that the qualitative spatial patterns for Macro-CXCL5, Tem-GZMK, and EC-FLT1 are preserved. (D) Sensitivity analyses for the association between SPP1 and disease-free survival across predefined subgroups, including the full cohort, exclusion of TRG 3, exclusion of TRG 0, and the residual-cancer-only subset.

Supplementary Figure 12

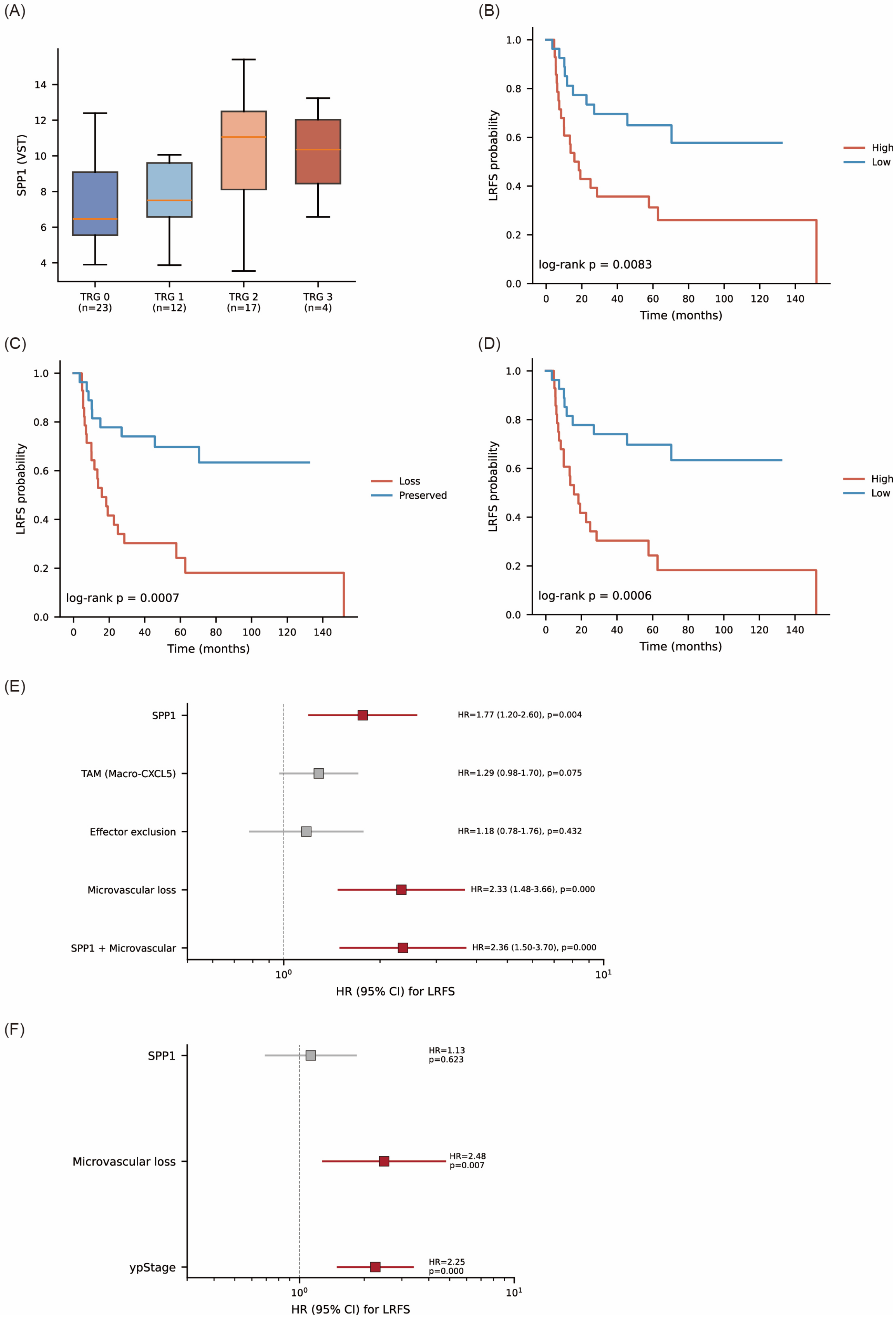

**Supplementary Figure 12. Locoregional recurrence-free survival analysis as a secondary endpoint.**

(A) Core-level SPP1 expression by tumor regression grade (TRG) group. (B) Kaplan–Meier curves for locoregional recurrence-free survival (LRFS) stratified by median-split SPP1 expression. (C) Kaplan–Meier curves for LRFS stratified by median-split microvascular-loss score. (D) Kaplan–Meier curves for LRFS stratified by the combined SPP1 + microvascular-loss score. Median splits were used for descriptive visualization only. (E) Univariable Cox proportional hazards models for LRFS using continuous z-scored resistant-niche features, including SPP1, TAM (Macro-CXCL5), effector exclusion, microvascular loss, and the combined SPP1 + microvascular-loss score. Hazard ratios are shown with 95% confidence intervals. (F) Multivariable Cox proportional hazards model for LRFS including SPP1, microvascular loss, and ypStage.

Supplementary Figure 13

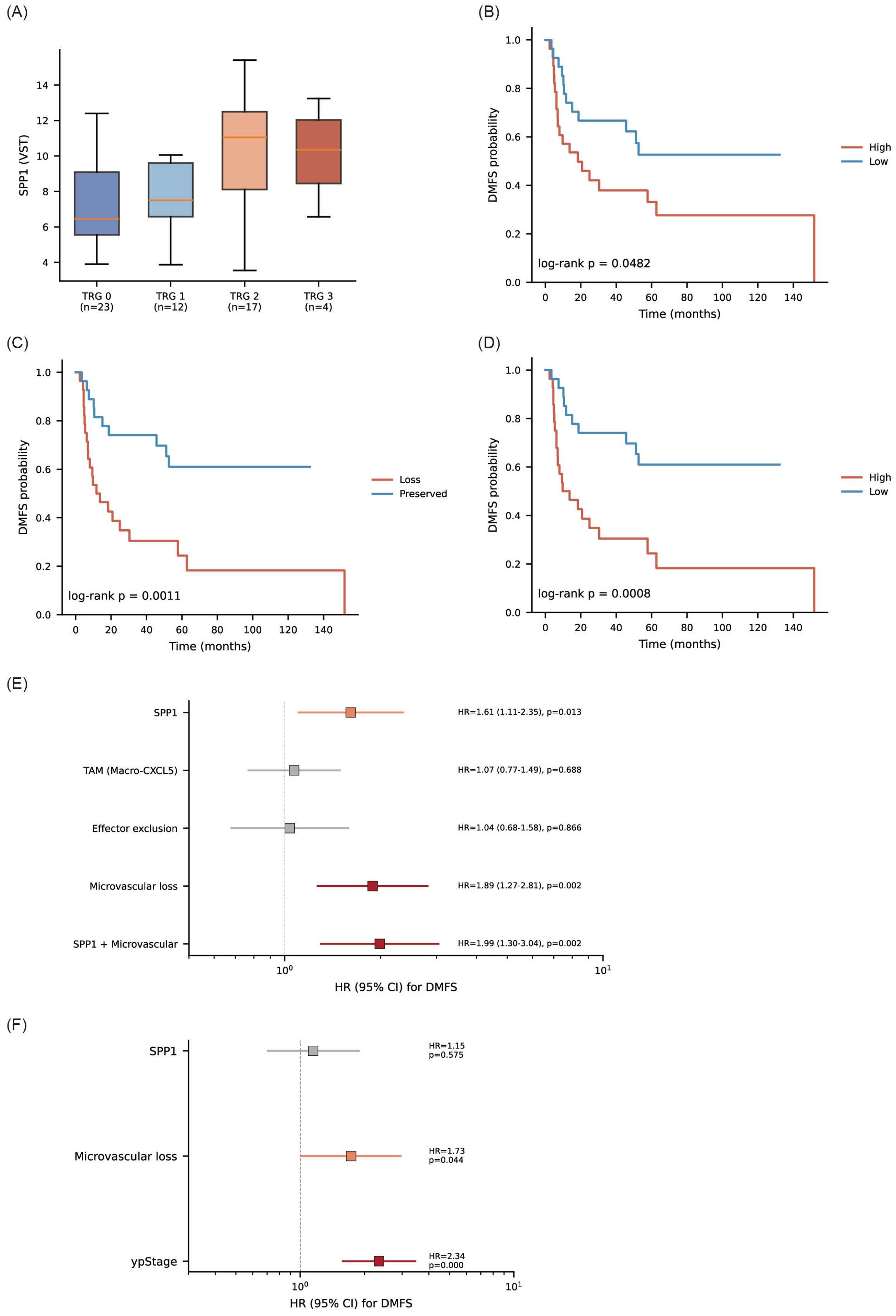

**Supplementary Figure 13. Distant metastasis-free survival analysis as a secondary endpoint.**

(A) Core-level SPP1 expression by tumor regression grade (TRG) group. (B) Kaplan–Meier curves for distant metastasis-free survival (DMFS) stratified by median-split SPP1 expression. (C) Kaplan–Meier curves for DMFS stratified by median-split microvascular-loss score. (D) Kaplan–Meier curves for DMFS stratified by the combined SPP1 + microvascular-loss score. Median splits were used for descriptive visualization only. (E) Univariable Cox proportional hazards models for DMFS using continuous z-scored resistant-niche features, including SPP1, TAM (Macro-CXCL5), effector exclusion, microvascular loss, and the combined SPP1 + microvascular-loss score. Hazard ratios are shown with 95% confidence intervals. (F) Multivariable Cox proportional hazards model for DMFS including SPP1, microvascular loss, and ypStage.

Supplementary Figure 14

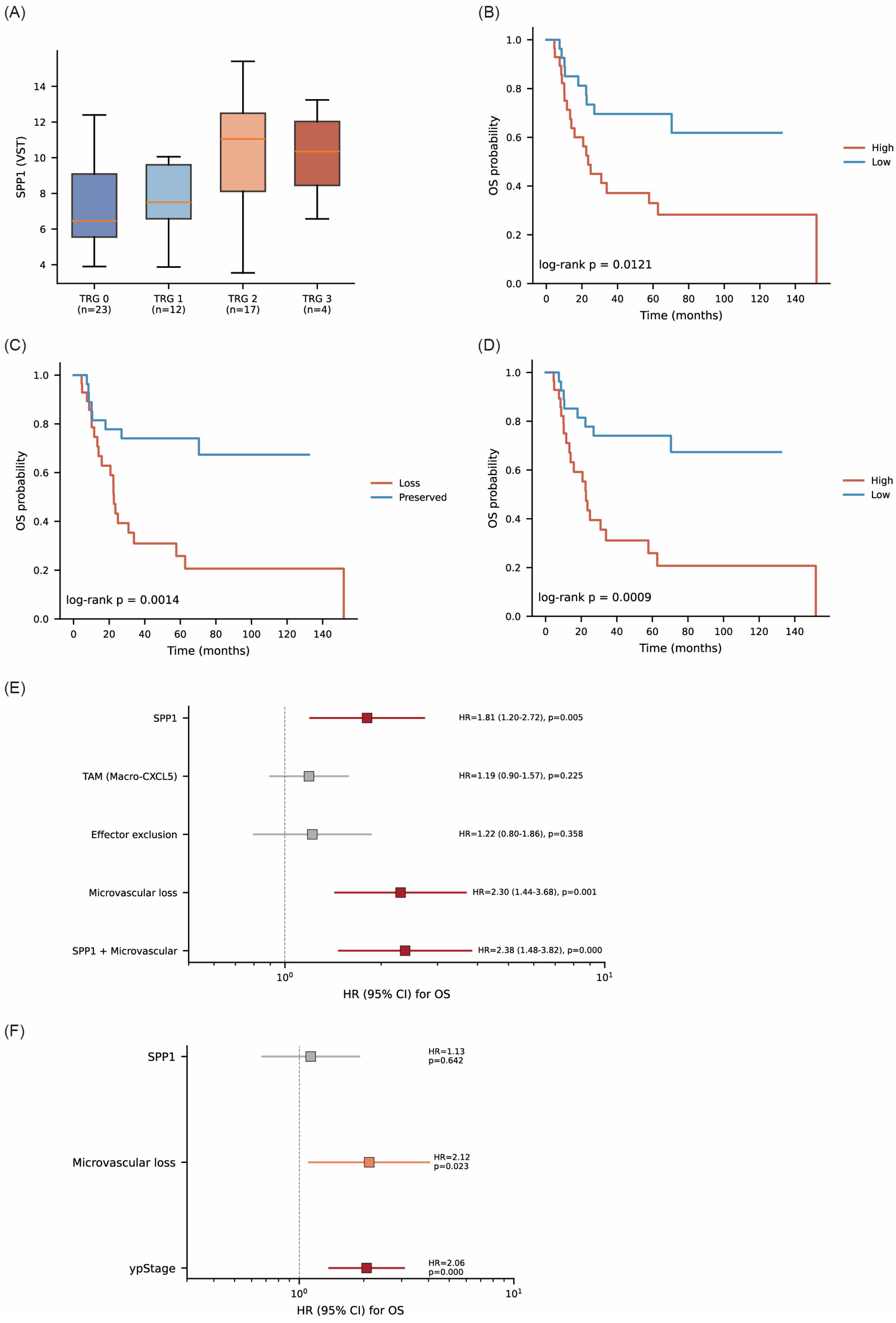

**Supplementary Figure 14. Overall survival analysis as an exploratory secondary endpoint.**

(A) Core-level SPP1 expression by tumor regression grade (TRG) group. (B) Kaplan–Meier curves for overall survival (OS) stratified by median-split SPP1 expression. (C) Kaplan–Meier curves for OS stratified by median-split microvascular-loss score. (D) Kaplan–Meier curves for OS stratified by the combined SPP1 + microvascular-loss score. Median splits were used for descriptive visualization only. (E) Univariable Cox proportional hazards models for OS using continuous z-scored resistant-niche features, including SPP1, TAM (Macro-CXCL5), effector exclusion, microvascular loss, and the combined SPP1 + microvascular-loss score. Hazard ratios are shown with 95% confidence intervals. (F) Multivariable Cox proportional hazards model for OS including SPP1, microvascular loss, and ypStage.

Supplementary Figure 15

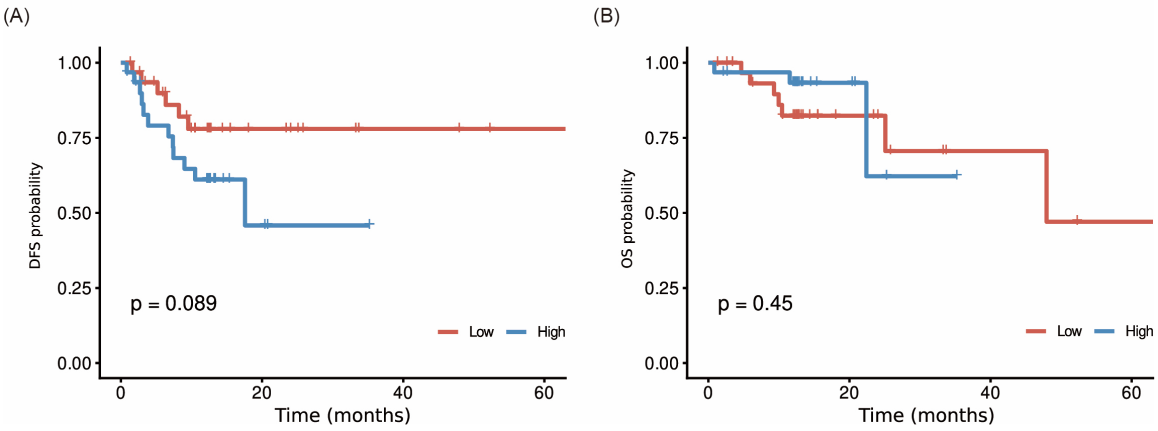


**Supplementary Figure 15. TCGA-ESCA ESCC cohort as orthogonal supportive clinical context for SPP1.**

(A) Kaplan–Meier curves for disease-free survival (DFS) in the TCGA-ESCA ESCC subset stratified by median-split SPP1 expression. Higher SPP1 expression showed a directionally unfavorable DFS trend. (B) Kaplan–Meier curves for overall survival (OS) in the TCGA-ESCA ESCC subset stratified by median-split SPP1 expression. No significant association with OS was observed.

Because TCGA-ESCA is composed predominantly of treatment-naïve resection specimens rather than post-CCRT residual disease, these analyses are presented as orthogonal supportive context rather than direct validation of the post-CCRT residual ESCC state.

### Supplementary Tables

**Supplementary Table S1**. Baseline clinicopathologic characteristics of the Visium cohort.

**Supplementary Table S2**. Top 20 SPP1-containing ligand–receptor pairs.

**Supplementary Table S3**. Top 30 ITGB1-containing ligand–receptor pairs.
